# Comparative genomics of zooxanthellate and azooxanthellate *Palythoa* (Cnidaria: Zoantharia) provides insights into symbiosis evolution and the origin of palytoxin

**DOI:** 10.64898/2026.04.08.717340

**Authors:** Yuki Yoshioka, Eiichi Shoguchi, Chiu Yi-Ling, Mayumi Kawamitsu, James Davis Reimer, Hiroshi Yamashita

## Abstract

**Background:** Zoantharians (Cnidaria) are globally distributed from shallow reefs to the deep sea, yet their genomic resources remain scarce. Within the zoantharian genus *Palythoa*, most species harbor photosymbiotic dinoflagellates, whereas some have secondarily lost this symbiosis, allowing genomic changes associated with symbiotic status to be examined. *Palythoa* is also the original source of palytoxin, one of the most potent marine toxins known, but whether it is produced by the host or by associated microorganisms remains unresolved. Here we generated high-quality genome assemblies of two zooxanthellate *Palythoa* species and compared five *Palythoa* genomes spanning both symbiotic states with those of other anthozoans.

**Results:** Across anthozoans, genome size scaled with transposable element content, particularly retrotransposons. The azooxanthellate lineage carries substitutions in conserved domains of the symbiosis-related proteins LePin and NPC2 and shows rapid evolution of the lectin gene *CLEC4A*, whereas NF-κB and V-type H^+^-ATPase remain conserved. *Palythoa* genomes showed no lineage-specific expansion of polyketide synthase (PKS) genes; cnidarian PKS genes were restricted to *P. caribaeorum* and *P.* sp. sakurajimensis (clade III). In contrast, 27 bacterial PKS sequences were recovered from metatranscriptomes and a metagenome-assembled genome across species, including an azooxanthellate species, indicating that they derive from associated bacterial rather than Symbiodiniaceae.

**Conclusions:** These genomes provide a comparative framework for zoantharians and reveal genomic changes associated with symbiotic status. We propose that palytoxin originates from associated bacteria rather than from the host, a hypothesis that can be tested by identifying the producing bacteria and their biosynthetic gene clusters.

## Introduction

Zoantharians (Cnidaria: Anthozoa: Hexacorallia) are globally distributed and occupy diverse habitats ranging from shallow coastal reefs to the deep sea [1, 2]. Most zoantharian species incorporate sand grains and detrital particles into their tissues, a distinctive trait thought to reinforce their body structure [3, 4]. Zoantharians are divided into two suborders. Macrocnemina species span a broad depth range and frequently engage in symbioses with a variety of marine organisms [5], whereas Brachycnemina are predominantly shallow-water taxa, most of which harbor photosynthetic dinoflagellates of the family Symbiodiniaceae [6]. Within Brachycnemina, *Palythoa* and *Zoanthus* are among the most abundant and conspicuous genera on coral reefs, where *Palythoa* species can exert strong ecological influences [7, 8] and, in some cases, produce palytoxin and/or palytoxin-like molecules (42-hydroxy-palytoxin) [9–12]. Although genome assemblies are now available for multiple actiniarian and scleractinian species [13], greatly advancing our understanding of anthozoan genome evolution [14–17], high-quality genomic resources and genomic studies for zoantharians remain comparatively sparse [18–20], limiting broader evolutionary inferences across Hexacorallia.

Within *Palythoa*, most species harbor Symbiodiniaceae, whereas *P. mizigama* and *P. umbrosa* inhabit low-light reef caves and lack Symbiodiniaceae symbionts, making them azooxanthellate [7]. Because symbiosis with Symbiodiniaceae likely originated in the common ancestor of Brachycnemina [21, 22], these species are thought to have secondarily lost algal symbiosis, making *Palythoa* a valuable system for comparing closely related zooxanthellate and azooxanthellate species. Our previous study using draft genomes of these two species revealed genomic features potentially associated with life in reef caves and the loss of algal symbiosis [20]. However, without genomes of zooxanthellate congeners, it remains unclear which of these features are shared across *Palythoa* and which are specific to azooxanthellate lineages.

Palytoxin, first isolated from *Palythoa toxica* [23], is one of the most potent marine toxins known, with a reported median lethal dose (LD_50_) of 25–450 ng/kg in several animal models [24]. Its exceptionally large and complex structure [25, 26] suggests biosynthesis by polyketide synthases (PKSs), in which ketosynthase (KS) domains serve as the core catalytic components [27, 28]. Complex polyketides in marine animals can be synthesized either by the animals themselves via PKS and animal fatty acid synthase-like PKS (AFPK) genes [29–31] or by bacterial symbionts [32–34]. Whether palytoxin in *Palythoa* is synthesized by the host or by associated bacteria therefore remains unresolved. Although some KS domain-containing genes were identified in draft genomes of *P. mizigama* and *P. umbrosa* [20], PKS genes have received little attention in other cnidarians, and no comparative survey within cnidarians has been conducted.

Here we present high-quality whole-genome assemblies and gene models for two zooxanthellate *Palythoa* species, *Palythoa* cf. *toxica* (Ptox) and *Palythoa* sp. sakurajimensis (Psak), together with curated genomes of the azooxanthellate *P. mizigama* (Pmiz) and *P. umbrosa* (Pumb) (Fig. 1a–c). Combined with a chromosome-level genome assembly of *Palythoa caribaeorum* (Pcar) from the Wellcome Sanger Institute, we performed comparative genomic analyses to (i) characterize genome evolution in zoantharians, (ii) identify genomic changes associated with symbiotic status by comparing zooxanthellate and azooxanthellate species, and (iii) evaluate the origin of palytoxin by surveying KS domain-containing genes in host genomes and associated microbes. This study provides genomic resources and candidate genes that establish a framework for future studies of symbiosis evolution and the origin of palytoxin in *Palythoa*.

**Fig. 1.**
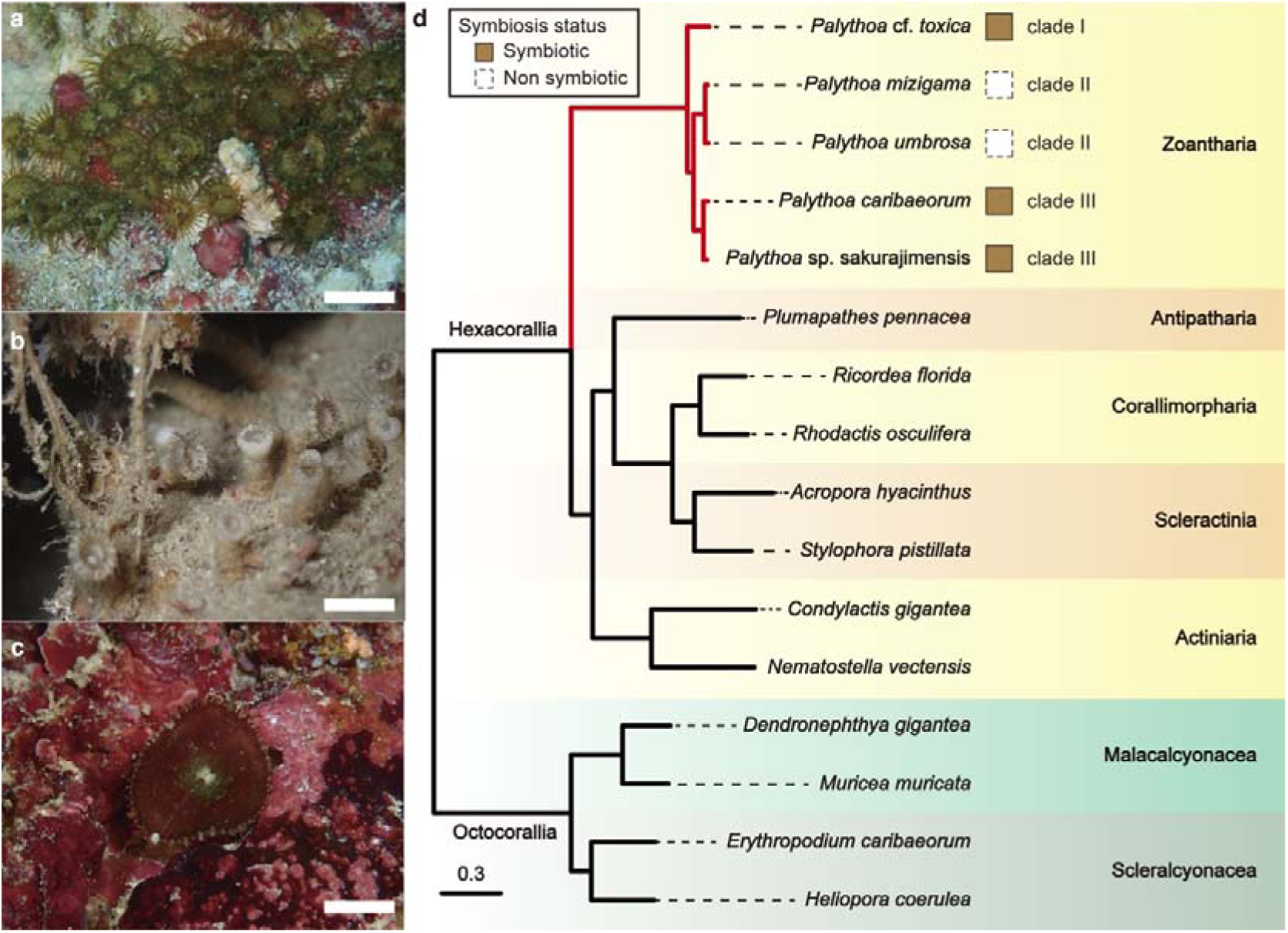
*Palythoa* species analyzed in this study and their phylogenetic position among anthozoans. **(a–c)** Colonies of **(a)** *Palythoa* cf. *toxica*, **(b)** *Palythoa umbrosa*, and **(c)** *Palythoa* sp. sakurajimensis (Photo credits: JDR). White scale bars indicate approximately 1 cm, 1 cm, and 2 cm, respectively. **(d)** Maximum-likelihood molecular phylogeny of anthozoans inferred from 1,887 single-copy orthologs (942,596 amino acid positions), with octocorals as the outgroup. All nodes received 100% bootstrap support (1,000 replicates). The scale bar indicates expected substitutions per site. Boxes indicate symbiotic status of *Palythoa* species with Symbiodiniaceae (filled, symbiotic; dashed, non-symbiotic), and clade numbers follow the main text. Symbiotic status is not shown for other species because it could not be confirmed for the sequenced samples.

## Results and Discussion

### High-quality genome assemblies and gene models of *Palythoa* species

We generated 50 Gb and 66 Gb of PacBio HiFi reads for Psak and Ptox, with mean read lengths of 5,433 bp and 5,599 bp, respectively (Table S1). The resulting draft genome assemblies spanned 880 Mb for Psak and 649 Mb for Ptox, with N50 values of 12.2 Mb and 0.82 Mb, respectively (Table 1; Fig. S1). Genome size estimates based on read depth were 819.3 Mb for Psak and 624.8 Mb for Ptox, indicating that the assemblies closely approximated the expected genome sizes [18]. In addition to these newly assembled genomes, we curated previously published genome assemblies of the azooxanthellate *Palythoa* species Pmiz and Pumb by removing contigs likely derived from contaminants (Fig. S2). BUSCO completeness scores for Psak, Ptox, and Pcar exceeded 90%, with low duplication rates (≤ 3%) (Table 1).

**Table 1.** Statistics for genome assembly and gene models.

Based on transcriptomic evidence and homology with metazoan orthologs, we predicted 20,594 protein-coding genes in Psak, 44,006 in Ptox, and 18,823 in Pcar (Table 1). Gene models for Pumb were re-predicted with transcriptomic evidence (22,498 genes), and those for Pmiz were re-predicted with metazoan protein evidence only because no RNA-seq data were available (27,878 genes) (Table 1). BUSCO completeness of the gene models was 86.2% (1.6% duplicates) for Psak, 75.0% (2.2% duplicates) for Ptox, 86.2% (1.8% duplicates) for Pcar, 77.8% (1.2% duplicates) for Pumb, and 78.9% (1.8% duplicates) for Pmiz (Table 1).

Complete mitochondrial genomes were also assembled for Psak (21,100 bp) and Ptox (21,227 bp), each encoding 13 protein-coding genes (Fig. S3). A previously reported novel mitochondrial ORF in *Palythoa tuberculosa* (Ptub) [35] was not detected in either species, supporting the hypothesis that the ORF was inserted in the common ancestor of Ptub and Pcar, rather than in the common ancestor of *Palythoa* clade III [35].

### Molecular phylogeny and genome-wide comparison among anthozoans

Using 1,887 single-copy orthologs, we reconstructed a maximum likelihood phylogeny of anthozoans (Fig. 1d). Consistent with analyses based on ultra-conserved elements [36], zoantharians were recovered as the sister group to a clade comprising Actiniaria, Antipatharia, Corallimorpharia, and Scleractinia. Within *Palythoa*, phylogenetic relationships inferred from nuclear single-copy orthologs were identical to those obtained using the mitochondrial *rrnL* marker, resolving three distinct clades: clade I (Ptox), clade II (azooxanthellate Pmiz and Pumb), and clade III (Psak and Pcar) (Fig. S4). This topology is congruent with previous phylogenetic reconstructions of zoantharians (e.g., Swain [22]).

Genome size varied substantially among anthozoans, including within *Palythoa* (Tables 1 and S2). Although more than 100 anthozoan genome assemblies are now publicly available [13], comparative analyses of genome-wide features across anthozoans remain limited. To place zoantharian genomes in this broader context, we examined how assembly size relates to genomic features across selected anthozoan species. A previous study suggested a correlation between assembly size and transposable element (TE) content in zoantharians [18], and we found that this relationship holds across anthozoans (Fig. 2). Assembly size was strongly correlated with total length of interspersed repeats (*r* = 0.968, *P* < 0.01) and moderately correlated with mean gene length (*r* = 0.700, *P* < 0.01) (Fig. 2). In contrast, neither protein-coding gene number nor mean CDS length showed a significant correlation with genome size. Non-TE sequence length (assembly size minus interspersed repeats) was also associated with assembly size, as was TE length, after phylogenetic correction (PGLS, λ = 0, *r* = 0.943–0.968, *P* < 0.01; Fig. S5a, b), indicating that genome expansion involved non-TE as well as TE sequences. Moreover, the TE proportion increased with assembly size (*r* = 0.617, *P* < 0.05), with a significant contribution from class I elements (retrotransposons; *r* = 0.593, *P* < 0.05) and a marginal contribution from unclassified elements (*r* = 0.498, *P* = 0.050), but not from class II elements (Fig. S5c–f). Thus, genome size variation in anthozoans is driven by the expansion of non-coding sequences, mainly retrotransposons together with introns and intergenic regions, as observed in other animals [37, 38].

**Fig. 2.**
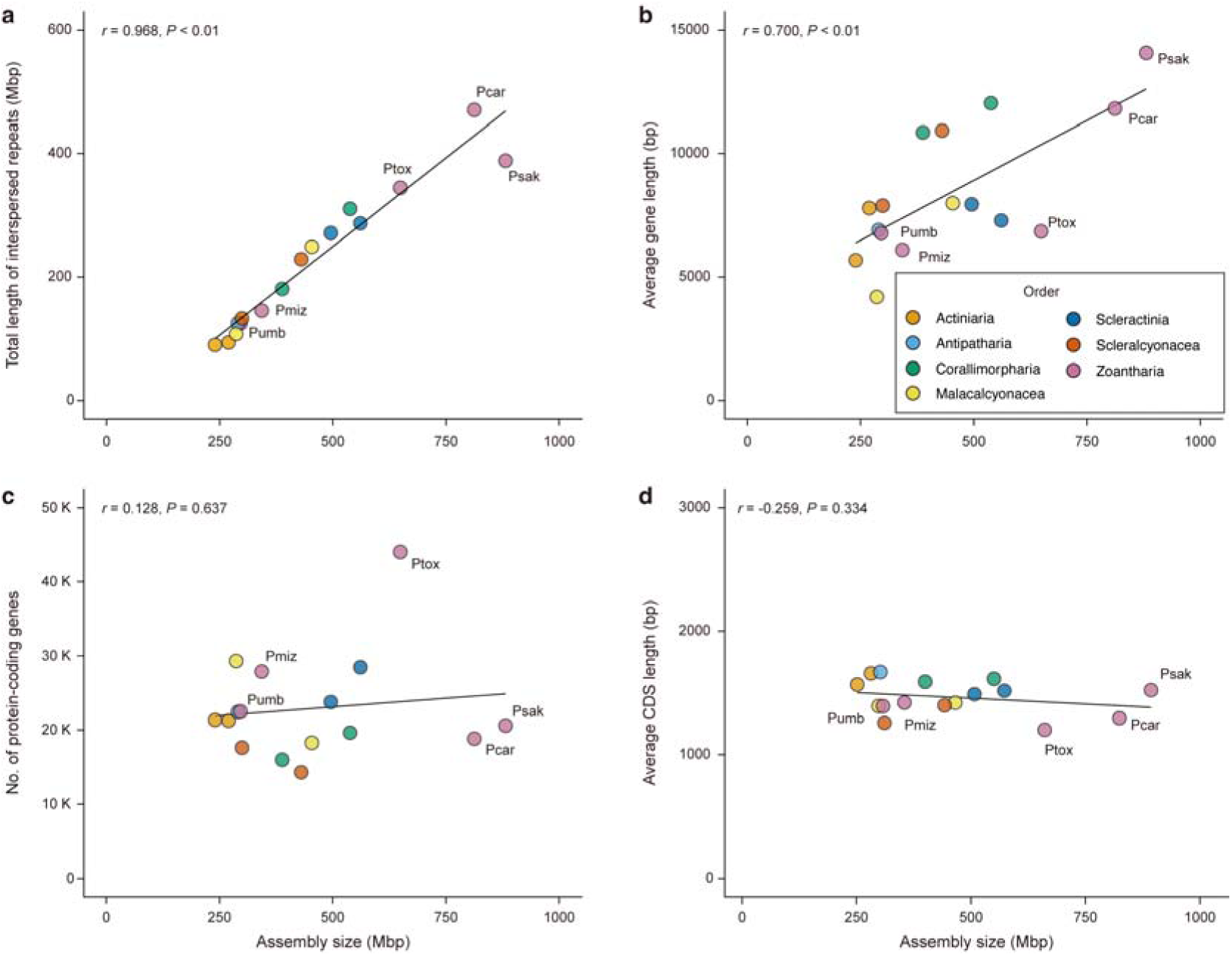
Relationships between assembly size and genome features across anthozoans. **(a)** Total length of transposable elements (Mb), **(b)** mean gene length, **(c)** number of protein-coding genes, and **(d)** mean CDS length, each plotted against assembly size. Each point represents one species, colored by order as indicated in (**b**). Lines indicate linear regressions, and *r* and *P* indicate Pearson’s correlation coefficients and their significance. PGLS results are shown in Fig. S5, and the underlying values are listed in Table S15.

### Transposable element activity and gene duplication in *Palythoa*

To infer the history of TE activity, Kimura substitution levels were calculated for all TE copies in each genome (Fig. S6). Lower Kimura distances indicate more recent transposition, whereas higher values reflect ancient transposition events. Clade I species (Ptox) displayed three pronounced peaks in Kimura distances, clade II species (Pmiz and Pumb) showed a diffuse pattern without clear peaks, and clade III (Psak and Pcar) exhibited a gradual decline with a peak at low divergence levels. Relatively high numbers of recent TE copies were commonly detected in all *Palythoa* clades, suggesting that genome size increases with TE amplification occurred relatively recently. TE-driven genome expansion has been shown to generate genomic diversity in *Hydra* [39], and a similar process may contribute to genomic diversity in *Palythoa* and perhaps other zoantharians.

Gene duplication also contributed to genome expansion, although to a lesser extent than TE insertion. With the exception of Ptox, protein-coding gene numbers were relatively uniform among anthozoans (Table S3). Psak, Pumb, Pmiz, and Pcar harbored 22,448 genes on average, whereas Ptox encoded 44,006 genes—nearly twice the average (Table 1). Furthermore, the proportion of intron-containing genes was markedly lower in Ptox (53.2%) than in other *Palythoa* species (69.8–78.9%; Table 1), suggesting extensive duplication of intron-less genes. Analysis of duplication mode revealed that dispersed duplications predominated (49–60%) in all *Palythoa* genomes, with Ptox exhibiting approximately twice as many dispersed duplications as other species (Fig. 3a). The wide spread of dispersed duplicates has been considered to be due to mechanisms mediated by transposons [40]. Most duplicated gene pairs showed *Ks* values below the median *Ks* of single-copy orthologs between *Palythoa* species, calculated from orthologs that are single-copy across hexacorals (Fig. 3b), indicating that the majority of duplication events occurred after *Palythoa* diversification.

**Fig. 3.**
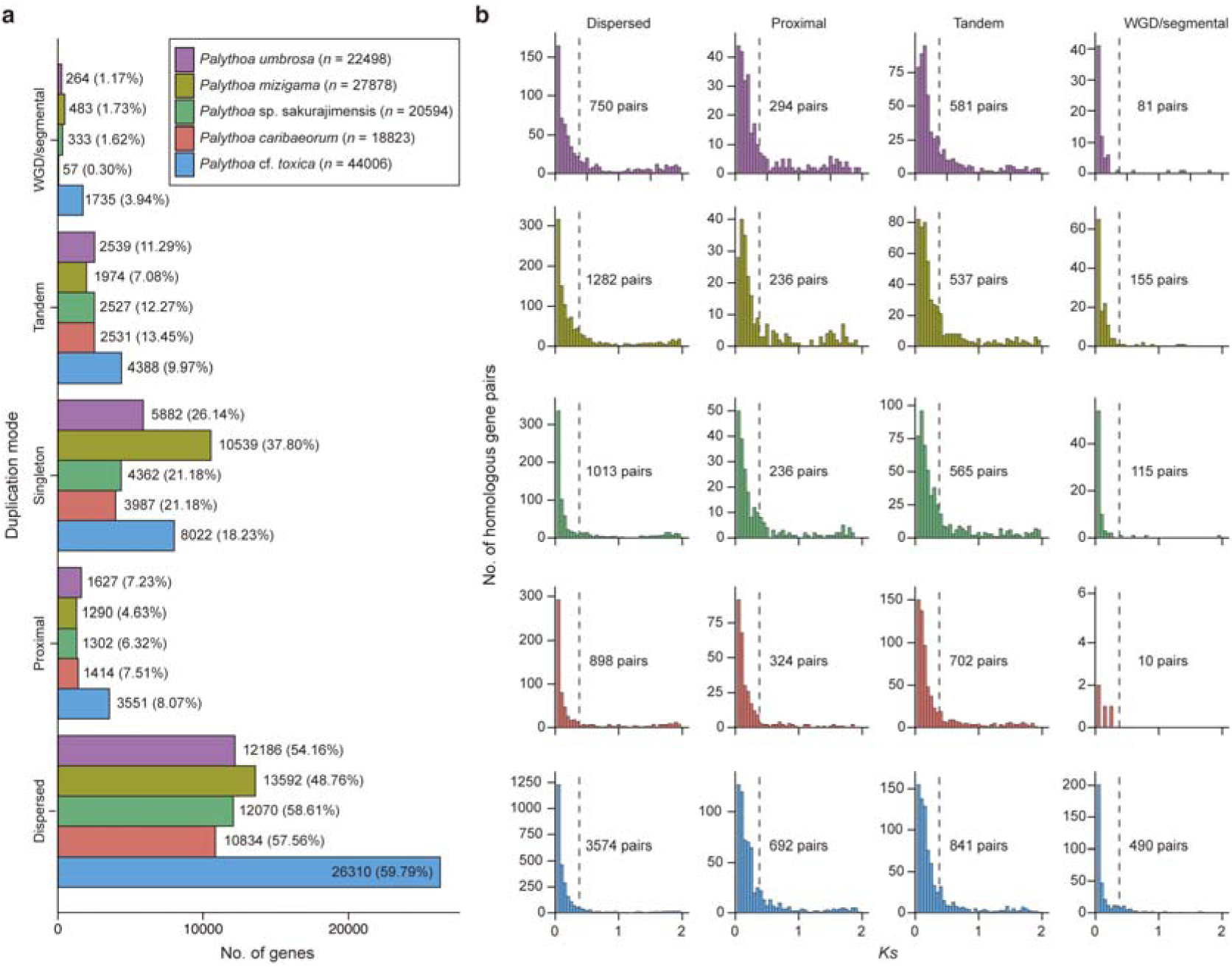
Gene duplication modes in *Palythoa*. **(a)** Number of genes in each duplication mode (dispersed, proximal, tandem, WGD/segmental, and singleton) classified by MCScanX. Numbers in parentheses indicate the percentage of all genes, and *n* indicates the number of predicted genes in each genome. **(b)** *Ks* distributions of paralogous gene pairs in each duplication mode. Dashed lines indicate the median *Ks* between *Palythoa* species, calculated from orthologs that are single-copy across hexacorals, and the number of gene pairs is shown in each panel. Colors indicate species as in (**a**).

### Gene family evolution in *Palythoa*

To investigate the molecular basis of zoantharian evolution, we examined gene family evolution across anthozoans. From 366,162 protein-coding genes, 93.6% were assigned to 26,532 orthogroups (OGs) (Table S4). Among these, 5,441 OGs were conserved across hexacorals, 342 OGs were specific to *Palythoa*, and 16 OGs were absent from *Palythoa* (Table S5).

Of the shared OGs, we analyzed the 4,944 OGs present in at least one octocoral. Using CAFE5, we identified 193 OGs that were significantly expanded in the common ancestor of *Palythoa* (FDR < 0.05; Fig. 4a; Table S6). These families were enriched in functions related to ion transport, including calcium channel activity, and to extracellular matrix (ECM) interactions, including glycosaminoglycan and heparin binding and metallopeptidase activity (Fig. 4b). These functions may be linked to a distinctive trait of zoantharians. Most zoantharians, particularly *Palythoa*, incorporate sand and detritus into their tissues to reinforce structural integrity [3]. As other anthozoans either secrete skeletons or maintain soft tissue without persistent foreign inclusions, incorporation of those particles likely requires specialized cellular processes. In *Hydra*, the mesoglea is an ECM containing collagens, laminin, and proteoglycans, and both its remodeling by matrix metalloproteinases and its interactions with cells, mediated in part by integrins, are required for morphogenesis and the maintenance of differentiated cells [41, 42]. The expansion of ECM-related gene families in *Palythoa* may therefore reflect the evolution of cellular machinery that actively integrates sediment into the mesoglea.

**Fig. 4.**
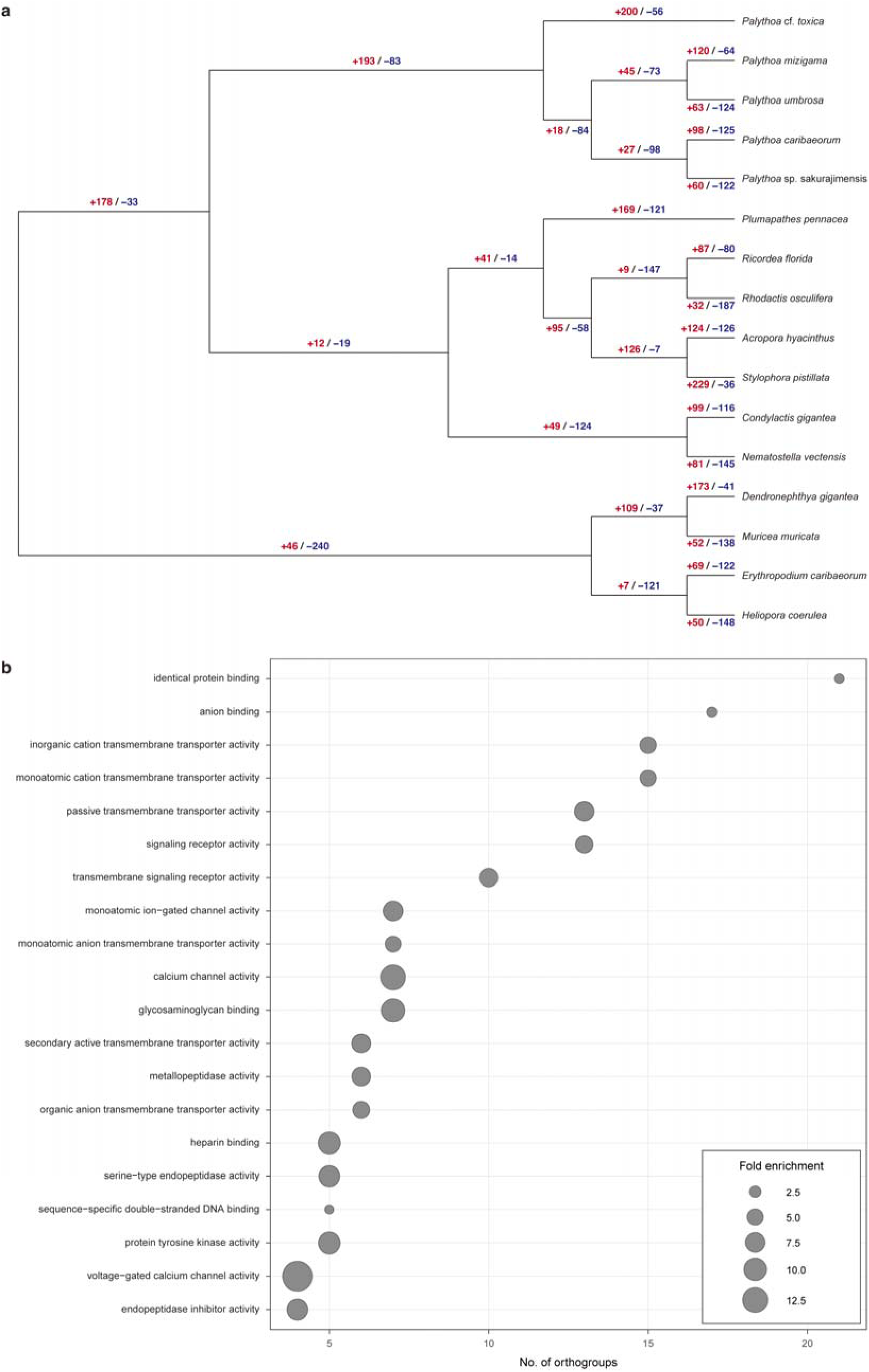
Gene family evolution across anthozoans and functions of families expanded in *Palythoa*. **(a)** Numbers of orthogroups (OGs) that significantly expanded (red) or contracted (blue) on each branch, inferred with CAFE5 (FDR < 0.05). **(b)** GO enrichment (Molecular Function) of the 193 OGs that expanded in the common ancestor of *Palythoa*. The top 20 terms are shown. The x-axis indicates the number of OGs, and circle size indicates fold enrichment. GO terms were assigned to *P. caribaeorum* proteins with eggNOG-mapper. The 4,944 OGs tested by CAFE5 served as the background, and enrichment was assessed using the weight01 algorithm with Fisher’s exact test, considering only GO terms annotated to at least five OGs.

Among the 342 *Palythoa*-restricted OGs, 289 (85%) were present as single copies or fewer than two copies per species (Fig. S7), suggesting that duplication of *Palythoa*-specific genes has been largely suppressed following lineage divergence. Functional annotation revealed that the most abundant categories among these OGs were “signal transduction mechanisms”, “post translational modification, protein turnover, chaperones”, and “transcription”, although “poorly characterized proteins” predominated (Fig. S8; Table S7). This pattern is consistent with taxonomically restricted genes (TRGs), which often lack annotated homologs but can play important roles in development and lineage-specific adaptations [43].

Sixteen OGs were inferred to be lost in *Palythoa*; of these, 11 were poorly characterized (Table S8). The remaining OGs included genes involved in chemoreception, sterol desaturation, and amide metabolism. Importantly, genes essential for amino acid biosynthesis were retained, indicating that *Palythoa* species likely retain the amino acid biosynthetic pathways present in other anthozoans. Compared with our previous study based on fragmented genomes and uncurated transcriptomes [20], the present results differ in some details. Nevertheless, the loss of a gene homologous to spermatogenesis-associated protein 7 (*SPATA7*), involved in photoreceptor ciliary function in mammals [44], was consistently detected, supporting its genuine absence in *Palythoa*. In addition, the visual pigment-like receptor peropsin (*RRH*) was also inferred to be lost, suggesting partial loss of photoreceptor-related functions across *Palythoa*, irrespective of symbiotic status.

### Clade-specific evolutionary changes within *Palythoa*

We identified 890, 484, and 107 OGs exclusive to clade I (Ptox), clade II (Pmiz and Pumb), and clade III (Pcar and Psak), respectively, containing 3,006 (Ptox), 586 (Pumb), and 143 (Pcar) genes in the representative genomes, respectively (Table S9). The majority of these genes lacked functional annotation (i.e., no COG category assigned or assigned to category S, “function unknown”): 84.1% (2,529 genes) in clade I, 75.1% (440 genes) in clade II, and 83.9% (120 genes) in clade III. Notably, 9.4% of clade I-specific genes encoded proteins with a reverse transcriptase (RVT) domain (Fig. S9; Table S9), suggesting lineage-specific expansion, particularly in Ptox.

Although OGs exclusive to each clade arose independently, they often shared homologous Pfam domains. In particular, genes containing reverse transcriptase (RVT_1; PF00078) and integrase (rve; PF00665) domains, both characteristic of retrotransposons, were recurrently detected in clades I and II (Fig. S9; Table S9). This pattern likely reflects the independent proliferation of retrotransposon-derived genes in each lineage, some of which may represent TE-derived open reading frames that escaped repeat masking. Such retrotransposon activity may also have increased gene numbers through RNA-mediated duplication [45], consistent with the high proportion of intron-less genes in Ptox.

### Rapidly evolving genes in *Palythoa*

We also identified paralogous gene pairs showing signatures of rapid evolution (*Ka*/*Ks* > 1). Ptox exhibited a high number of rapidly evolving paralogous gene pairs (20 pairs), whereas other species showed fewer (5 pairs each) (Fig. S10; Table S10). Notably, tumor protein translationally-controlled 1 (*TPT1*: OG0001769) showed rapid evolution in clade III species, while C-type lectin domain family 4 member A (*CLEC4A*: OG0002049) evolved rapidly in clade II. *TPT1* is involved in cell growth and proliferation [46] and its rapid evolution might therefore be related to the rapid growth and ecological dominance of clade III species, such as Pcar, on coral reefs [47–50]. *CLEC4A* binds carbohydrates abundant on cell surfaces, such as mannose and N-acetylglucosamine (GlcNAc) [51], and shows broad glycan-binding specificity in humans [52]. Because even a few substitutions in the carbohydrate-recognition domain can shift sugar-binding specificity [53], rapid evolution of *CLEC4A* could either narrow its recognition to particular glycans or further broaden it. Such changes might facilitate association with specific bacteria in azooxanthellate clade II and contribute to adaptation to reef caves. Both hypotheses remain to be tested by functional analyses.

### Loss of fluorescent proteins and divergence of symbiosis-related genes in azooxanthellate *Palythoa*

Amplicon sequencing of symbiotic algae revealed distinct Symbiodiniaceae associations: Psak was dominated by *Durusdinium* (74.2%), whereas Ptox predominantly harbored *Cladocopium* (98.3%) (Table S11). Although extensive examination of symbiotic algae across host clades was not performed, given that symbiosis with Symbiodiniaceae likely originated in the common ancestor of Brachycnemina [21, 22], transitions of algal partners likely occurred during *Palythoa* evolution. Alternatively, species in *Palythoa* clade III may exhibit greater flexibility in their symbiotic associations [54–56].

Genes encoding fluorescent protein homologs (OG0000497) were absent from clade II (Pmiz and Pumb). To trace the evolutionary history of this gene family, we examined microsynteny around the eight fluorescent protein homologs in the chromosome-level Pcar genome, which are distributed across three chromosomes and include two tandem clusters (g726–g727 and g742–g745) (Fig. S11). Around g496 and g726–g727, flanking genes were retained on single contigs in both Pmiz and Pumb, but no fluorescent protein homolog was present between them, and none was detected by BLASTN or HMM searches of six-frame translations of either assembly. For the other regions, the corresponding loci were interrupted by contig ends in the azooxanthellate assemblies. Although Ptox possesses 10 homologs, none was located at the corresponding loci (Fig. S11a, b). These results suggest that fluorescent protein homologs in *Palythoa* have undergone repeated duplication and loss rather than being stably inherited at fixed genomic positions. The absence of fluorescent protein homologs in Pmiz and Pumb is consistent with the lack of detectable autofluorescence in these species [7]. Autofluorescence, mostly due to green fluorescent protein, is widespread in cnidarians and has been proposed to function in prey attraction [57, 58], algal symbiont attraction [59, 60], and fine-tuning of light microclimate [61, 62]. These functions depend on ambient light, and some of them also depend on algal symbionts, so they are likely to be of limited value for azooxanthellate species inhabiting low-light reef caves. Relaxed selection associated with low-light habitats and the loss of algal symbionts may have allowed the gene family to be lost entirely in the lineage leading to Pmiz and Pumb. These findings suggest a reorganization of light-dependent and feeding-related functions in azooxanthellate zoantharians, although targeted ecological and physiological studies will be required to clarify the underlying mechanisms.

A recent study demonstrated that LePin (lectin and Kazal protease inhibitor domains) is involved in the initiation of endosymbiosis in the octocoral *Xenia* sp. [63]. Although this gene is widely conserved among anthozoans, structural divergence has been suggested in the azooxanthellate sea anemone *Nematostella vectensis* [63]. In *Palythoa* species, some LePin homologs were identified in OG0000224 (Table S5). Some homologs retained the conserved domain architecture observed in *Xenia* (i.e., C-type lectin, H-type lectin, EGF-like, and Kazal domains), whereas others exhibited more diverse protein domain architectures (Fig. 5). Comparison of amino acid sequences that retained conserved domain architecture revealed five sites with substitutions specific to azooxanthellate lineages, four of which were located within annotated functional domains (H-type lectin, C-type lectin, and EGF-like domains) and were conserved among zooxanthellate *Palythoa* and *Xenia*. Although the functional consequences of these substitutions remain unclear, their localization within key domains suggests potential functional modification or reduction of LePin in azooxanthellate *Palythoa*.

**Fig. 5.**
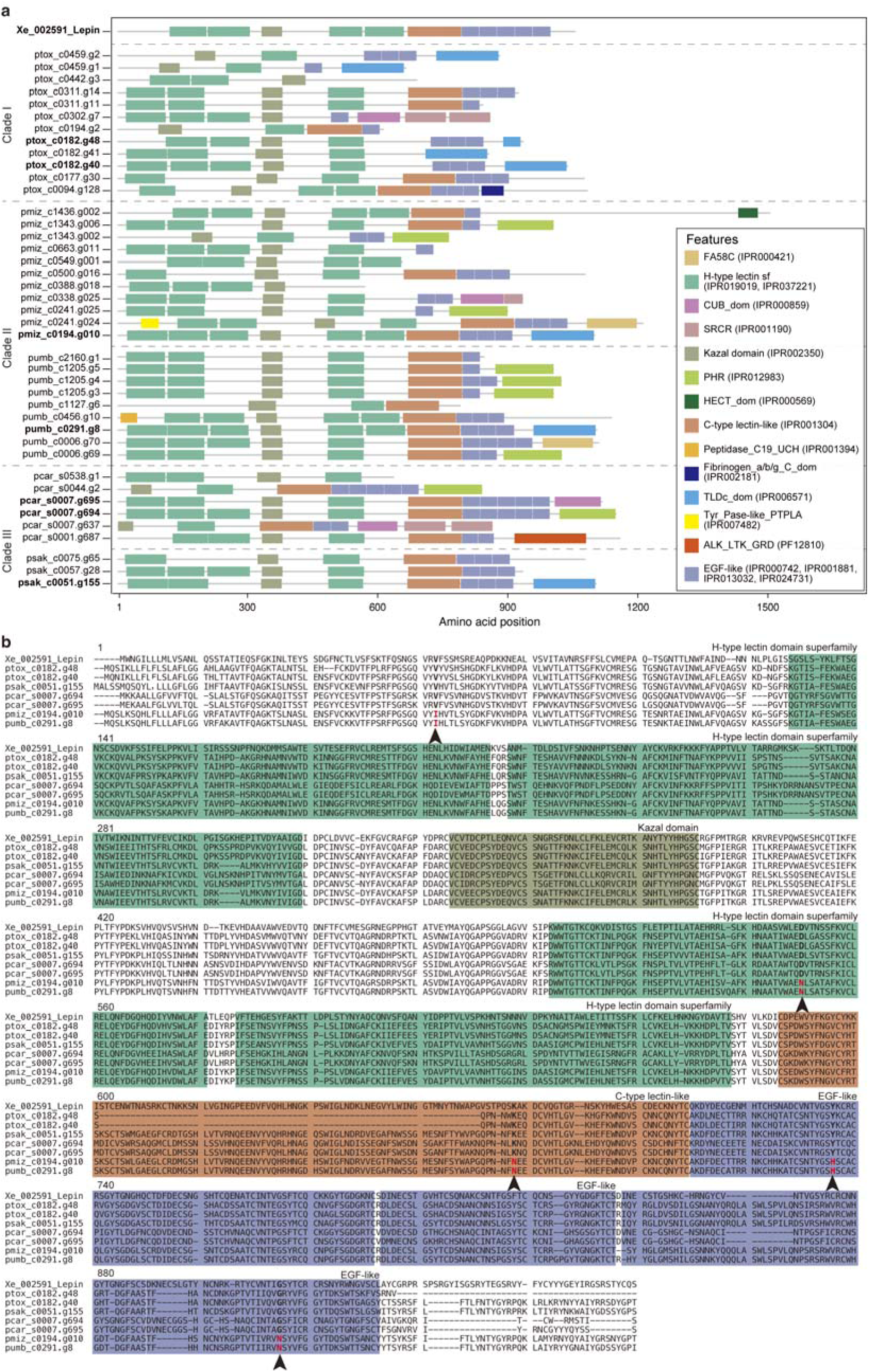
LePin homologs in *Palythoa* genomes. **(a)** Domain architectures of LePin homologs in *Palythoa* and *Xenia* sp. (Xe_002591). Colors indicate protein domains, with InterPro or Pfam entries in parentheses. Genes that retain the domain architecture of Xenia LePin are shown in bold. **(b)** Amino acid alignment of the homologs with the conserved domain architecture. Domain regions are colored as in (**a**). Arrowheads and red letters indicate substitutions specific to azooxanthellate *Palythoa*.

We further examined three proteins central to cnidarian-dinoflagellate symbiosis: the transcription factor NF-κB, which regulates host immune responses during symbiosis [64, 65], a proton pump V-type H^+^-ATPase (VHA), which is present in the host membrane surrounding the algal symbionts and acidifies the symbiosome to promote algal photosynthesis [66], and the sterol transporter Niemann-Pick type C2 (NPC2), which mediates sterol transfer between symbionts and hosts [67–69]. No azooxanthellate-specific substitutions were found in NF-κB (Fig. S12) or VHA (Fig. S13), consistent with their broad roles in immunity and cellular homeostasis across animals. In contrast, azooxanthellate-specific substitutions were found within conserved domain regions in two of the four NPC2 paralog groups (Fig. S14). Thus, lineage-specific changes in azooxanthellate *Palythoa* were detected in genes involved in symbiont recognition (LePin) and nutrient exchange (NPC2), whereas the broadly acting NF-κB and VHA remained conserved. These substitutions may reflect functional modification or relaxed selective constraint. However, because both azooxanthellate species belong to a single clade (clade II), we cannot determine whether they contributed to or resulted from the loss of symbiosis, or arose in the clade II ancestor for unrelated reasons; addressing this will require comparison with independently derived azooxanthellate lineages and experimental validation.

### Bacterial rather than host PKS genes as candidate sources of palytoxin

We identified 166 KS domain-containing gene candidates, of which 95 passed the score threshold of AFPK-Finder (Fig. 6). Molecular phylogenetic analysis grouped these into four clades: FAS, FAS II, cnidarian PKS, and bacterial PKS (Fig. 6). FAS genes, which are highly conserved among animals and are responsible for synthesizing saturated fatty acids from acetyl-CoA and malonyl-CoA, were present in all examined hexacorals except actiniarians, including all five *Palythoa* species. Most cnidarian PKS proteins contained a patched domain (IPR003392) in addition to the canonical KS, AT, DH, KR, and TE domains (Fig. S15). Within *Palythoa*, cnidarian PKS genes were detected only in the genomes of Pcar and Psak (Fig. 6). Microsynteny analysis showed that, in Ptox, the genes flanking the PKS locus were retained on a single contig but no PKS gene was present between them (Fig. S16a). In Pmiz and Pumb, the corresponding region was interrupted by contig end; however, no cnidarian PKS gene was detected anywhere in these assemblies by BLSATN or HMM searches of six-frame translations, indicating that this locus is restricted to Pcar and Psak among the species examined. Pcar carried two copies, and microsynteny suggested that the ancestral copy became a pseudogene, containing two premature stop codons, after duplication (Fig. S16). Notably, no lineage-specific PKS expansion was observed in any *Palythoa* genome.

**Fig. 6.**
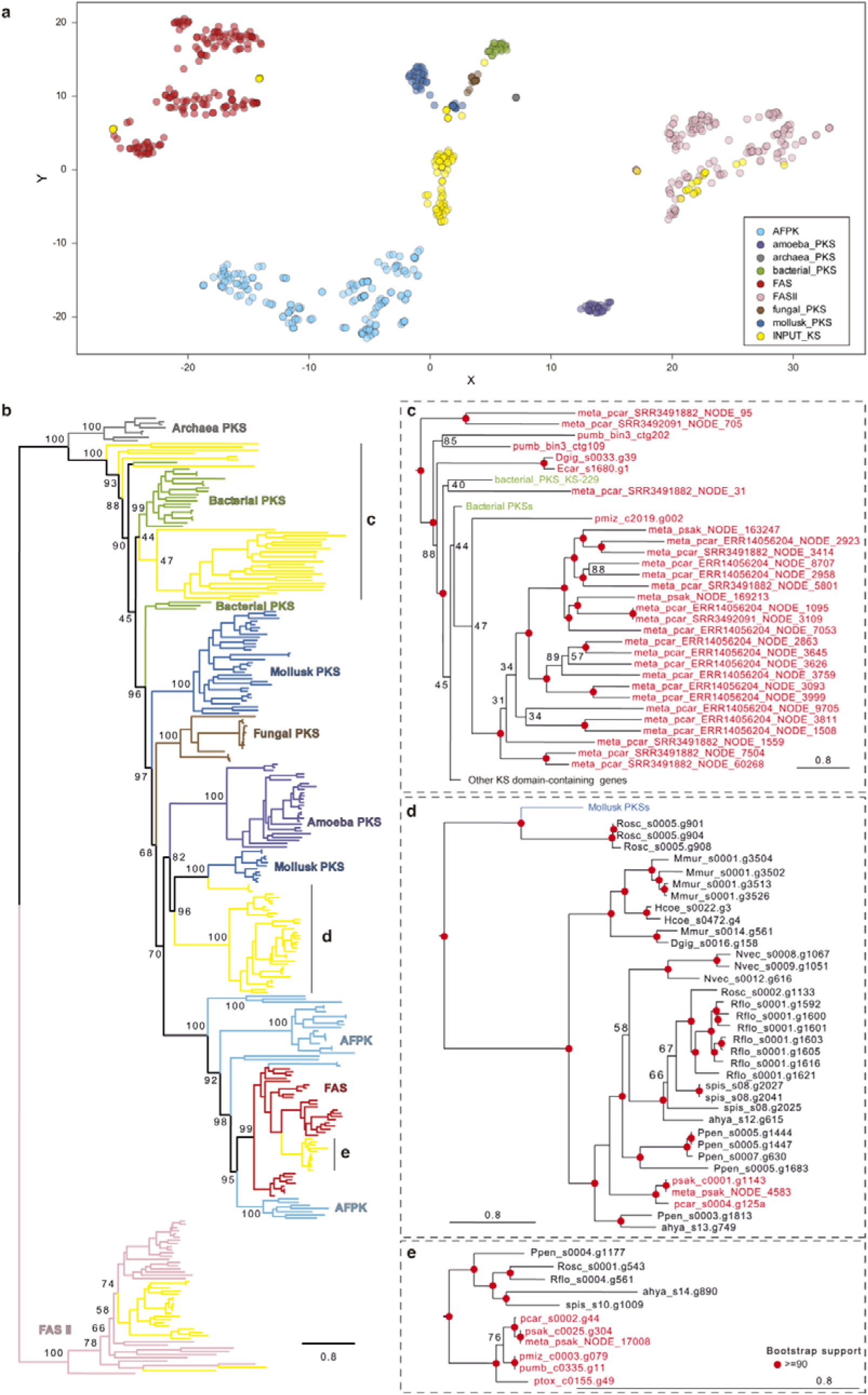
Phylogenetic distribution of ketosynthase (KS) domain-containing genes in anthozoans and associated microbes. **(a)** t-SNE plot generated by AFPK-Finder. Colors indicate clades defined by Lin et al. [31], and candidates identified in this study are shown in yellow. **(b)** Maximum likelihood tree of KS domains. KS sequences from non-anthozoan organisms were retrieved from the AFPK-Finder dataset. Colors indicate clades as in (**a**), and branches containing cnidarian sequences are shown in yellow. Bootstrap values (1,000 replicates) are shown for major nodes. **(c–e)** Enlarged views of the clades containing (**c**) bacterial PKSs, **(d)** cnidarian PKSs, and **(e)** FASs. *Palythoa* sequences are shown in red. “meta” and “bin” indicate sequences from metatranscriptome and metagenome-assembled genomes, respectively. Red circles indicate bootstrap values ≥ 90. Scale bars indicate substitutions per site. The alignment file is available in Zenodo, and domain architectures of cnidarian genes are shown in Fig. S15. PKS, polyketide synthase; FAS, fatty-acid synthase; AFPK, animal fatty-acid synthase-like polyketide synthase.

In contrast, bacterial PKS sequences were recurrently detected in the microbial fraction of *Palythoa* samples. We recovered 25 such sequences from metatranscriptomes of Pcar and Psak and two from a MAG assigned to Campylobacterota, reconstructed from Pumb reads (Fig. 6; Tables S12–S14). Their recurrent detection in three independent Pcar datasets and in Psak, together with the bacterial MAG from the azooxanthellate Pumb, indicates that PKS-bearing bacteria are consistently associated with *Palythoa* and that these genes are not derived from Symbiodiniaceae. No bacterial PKS sequences were recovered from Ptox and Pumb metatranscriptomes; however, these yielded far fewer predicted proteins that those of Pcar and Psak (approximately 24,000 vs. 83,000–288,000; Table S13), and Ptox RNA was partially degraded (Table S1), so this absence likely reflects limited sequencing depth. Only one bacterial-like PKS gene (pmiz_c2019.g002) was found in a *Palythoa* genome assembly. This gene contained three introns, indicating that it is integrated into the host genome rather than derived from bacterial contamination, and suggesting acquisition through horizontal gene transfer. Together, the absence of host PKS expansion, the absence of cnidarian PKS from Ptox, and the recurrent detection of expressed bacterial PKS genes in *Palythoa*-associated microbes suggest that associated bacteria, rather than host-encoded PKSs, are the most likely source of palytoxin.

### Conclusions

We present new genome assemblies of two zooxanthellate *Palythoa* species and curated assemblies of two azooxanthellate species, which, together with the chromosome-level genome of *P. caribaeorum*, provide a comparative genomic framework for zoantharians. Across anthozoans, genome size variation was closely associated with TE content, particularly retrotransposons. Gene families related to ion transport and extracellular matrix interactions expanded in the common ancestor of *Palythoa*, possibly in association with the incorporation of sand into the mesoglea. Comparisons between zooxanthellate and azooxanthellate species revealed genomic changes associated with symbiotic status. The azooxanthellate lineage has lost a fluorescent protein gene family that otherwise undergoes repeated duplication and loss, and shows lineage-specific substitutions in the symbiosis-related proteins LePin and NPC2 and rapid evolution of the lectin gene *CLEC4A*, whereas NF-κB and VHA remain conserved. Regarding the origin of palytoxin, *Palythoa* genomes showed no lineage-specific expansion of PKS genes, whereas bacterial PKS genes were recurrently detected and expressed in *Palythoa*-associated microbial communities. Based on these findings, we propose that palytoxin originates from associated bacteria rather than from the host. Testing this hypothesis will require identifying the producing bacteria and their biosynthetic gene clusters, relating their abundance and expression to palytoxin content across species and colonies, and experimental approaches such as antibiotic treatment of host colonies or cultivation of candidate bacteria.

## Materials and methods

### Sample preparation, DNA extraction, and long read sequencing

Ptox and Psak were collected between September 16-17, 2023 from reefs around Iriomote Island, Okinawa, Japan, at depths of ∼20 m and 5 m, respectively, and maintained for approximately twenty months in aquaria at the Yaeyama Station, Fisheries Technology Institute (Okinawa, Japan). For DNA extraction, a single polyp was collected from each colony, cut into small pieces using a razor blade, and briefly dissociated with Trypsin/EDTA (T4049, Sigma-Aldrich) for 5 min at room temperature. After digestion, the suspension was centrifuged at 500 × *g* for 5 min, and the resulting pellet was washed with artificial seawater filtered through 0.22 µm membranes (FSW). High molecular weight DNA was extracted from the pellet using the MagAttract HMW DNA kit (Qiagen) following the manufacturer’s protocol. Sequencing libraries were prepared using the HiFi DNA protocol (Pacific Biosciences) for Psak and the HiFi ultra-low input DNA protocol for Ptox. Libraries were sequenced on the PacBio Revio system (Pacific Biosciences).

In addition, to improve previously reported gene models of Pumb, polyps of the species were also collected and processed in the same manner as the other two species from a specimen collected on September 16, 2023 from Nakano Beach, Iriomote Island at 5 m depth.

Polyps from the *Palythoa* colonies used for this work have been deposited in the University of the Ryukyus Fujukan Museum under specimen numbers XXXXX.

### RNA extraction and sequencing

Fresh polyps of Psak, Ptox, and Pumb were individually homogenized in TRIzol reagent (Thermo Fisher Scientific). Total RNA was extracted according to the manufacturer’s protocol and its quantity and quality were assessed using Qubit RNA Assay Kit (Thermo Fisher Scientific) and a TapeStation (Agilent). RNA-seq libraries were constructed using the NEBNext Ultra II Directional RNA Library Prep Kit for Illumina and sequenced on an Illumina NovaSeq X with 150-bp paired-end mode (Table S1).

### *De novo* genome assembly

HiFi reads potentially originating from non-cnidarian organisms were identified with FCS-GX v0.5.4 [70] and were removed from further steps. Mitochondrial genomes were first assembled from the filtered HiFi reads using the MitoHiFi pipeline v3.2.1 [71]. Reads mapping to mitochondrial genomes were subsequently removed prior to nuclear genome assembly.

Contaminant- and mitochondria-free HiFi reads were assembled using Hifiasm v0.23.0-r691 [72]. Primary contigs generated by Hifiasm were used for downstream analyses. Haplotigs were removed using a combination of the kmerDedup pipeline (https://github.com/xiekunwhy/kmerDedup) and purge_dups v1.2.5 [73]. Assembly polishing was performed using Inspector v1.3 [74] with filtered HiFi reads.

Assemblies were evaluated using BlobTools v1.1.1 [75]. Contigs classified as taxa other than “Cnidaria” or “no-hit” were considered as non-*Palythoa* sequences and excluded from further analyses. Genome completeness was assessed using BUSCO v6.0.0 [76] with the “metazoa_odb12” dataset (*n* = 672). Genome size was estimated using the backmap.pl v0.6 pipeline [77], in which filtered HiFi reads were mapped back to the draft assembly using minimap2 v2.26-r1175 [78] with the option (-ax map-hifi).

### Additional genomic datasets

For comparative analyses, the genome assembly of *Palythoa caribaeorum* (Pcar; chromosome-level, GCA_965234985) was retrieved from NCBI. Genome assemblies of Pumb and Pmiz (contig-level), previously reported in Yoshioka et al. [20], were also included and curated using BlobTools, with non-cnidarian contigs removed. In addition, genome assemblies generated using long-read sequencing (PacBio or Oxford Nanopore) from representative species across six anthozoan orders were included for comparative analyses (Tables S2 and S3).

### Identification of repetitive elements and divergence analysis

Repetitive elements were identified *de novo* using RepeatModeler v2.0.4 [79] with the option (-LTRStruct). Repeats identified *de novo* and known repeats from RepeatMasker database were combined and soft-masked using RepeatMasker v4.1.5 [80]. Transposable elements’ (TE) divergence and landscape analyses were performed using the RepeatMasker scripts “calcDivergenceFromAlign.pl” and “createRepeatLandscape.pl”. TEs were classified into nine categories: DNA transposons, LINEs, LTRs, PLEs, rolling circle (RC)/Helitrons, Retroposons, Satellites, SINEs, and unclassified elements. When we performed correlation analysis with genome size, interspersed repeats were further grouped into three: class I Retroelements (LINEs, SINEs, LTR, PLE and retroposons), class II DNA transposons (DNA transposons and RC/Helitrons) or Unclassified.

### Protein-coding gene annotation

RNA-seq reads were quality trimmed using fastp v1.0.1 [81] with a minimum read length of 50 bp. Filtered reads were assembled *de novo* using TRINITY v2.15.1 [82] with default parameters. To remove low-abundance transcripts, reads were mapped back to the transcriptome assembly, and only concordantly mapped reads were retained as in [83]. These reads were aligned to the genome assembly using HISAT2 v2.2.1 [84].

Gene prediction was performed using BRAKER pipeline v3.0.8 [85–87], with soft-masked genomes, RNA-seq alignments, and Metazoa OrthoDB v12 as input. Gene models were further refined using PASA v2.5.2 [88] with genome-guided transcriptome assemblies generated by TRINITY. Genes with CDS length greater than 99 bp were retained. Gene identifiers were assigned sequentially based on contig position (e.g., c0001.g001 is the first gene of contig c0001). Gene model completeness was assessed using BUSCO v6.0.0 with the “metazoa_odb12” dataset. RNA-seq data used for gene prediction are listed in Table S1.

### Orthogroup classification and gene duplication analysis

For each species, the longest CDS per gene was selected and translated into protein sequences using TransDecoder v5.7.1 (https://github.com/TransDecoder/), following Yoshioka et al. [89]. Orthogroups (OGs) were inferred using OrthoFinder v2.5.5 [90]. Duplicated genes were classified as dispersed, proximal, singleton, tandem, or whole genome duplication (WGD)/segmental using MCScanX v1.0.0 [40].

For gene family expansion and contraction analysis, we used Computational Analysis of gene Family Evolution (CAFE5) v5.1.0 [91]. Orthogroups in which any single species had more than 100 gene copies were removed, because such families cause CAFE’s parameter estimates to be non-informative. The maximum-likelihood species tree inferred from 1,887 concatenated single-copy orthologs was rooted with the octocorals using ape v5.8-1 [92] in R. The tree was then made ultrametric by penalized likelihood with “chronos()” (correlated-rate model, smoothing parameter λ = 1). A single error rate (ε) for gene counts was first estimated to allow for assembly and annotation errors, which could differ among genomes of different completeness. The base model was then refitted with this error distribution fixed. For each orthogroup, CAFE5 computes a family-wide *P* value for deviation from the fitted birth–death model. We corrected these *P* values for multiple testing with the Benjamini-Hochberg method. For each branch, we counted the significant orthogroups that expanded or contracted between the ancestral and descendant nodes reconstructed by CAFE5.

### Gene duplication history and molecular evolution

To examine gene duplication history, paralogous gene pairs were identified within each genome, and pairwise nonsynonymous (*Ka*) and synonymous (*Ks*) substitution rates were calculated following Mathers et al. [93]. All-against-all BLASTP searches were performed among genes within each duplication category. Paralog pairs were retained if they aligned over ≥150 amino acids with ≥30% identity, and for each gene only its closest paralog was kept. Because each pair was detected in both directions (e.g., A–B and B–A), only one of the two was retained. Finally, to avoid ambiguous assignments, pairs whose genes were not assigned to the same OG were excluded.

Protein alignments were generated using MAFFT [94] with the E-INS-i strategy, and corresponding codon alignments were obtained using PAL2NAL [95]. *Ka* and *Ks* values were calculated using KaKs_Calculator 2.0 [96] under the Yang-Nielsen model. Paralog pairs with *Ks*□<□0.01 or *Ks* >□2 were excluded. Median *Ks* values of single-copy orthologs among hexacorals were used to classify duplications as pre- or post-speciation events. Multiple testing corrections were applied using the Benjamini-Hochberg method. Gene pairs with *Ka*/*Ks*□>□1 and adjusted *P* < 0.05 were considered to be under positive selection.

### Molecular phylogenetic analyses

Nucleotide or protein sequences were aligned using MAFFT v7.525 with the option (--auto), and poorly aligned regions were trimmed using ClipKIT v2.2.4 [97]. Concatenated alignments were generated using PhyKIT v2.0.1 [98]. Maximum likelihood phylogenies were inferred using IQ-TREE v2.2.2.6 [99] with ModelFinder [100] (-m MFP or -m MFP+MERGE) and UFBoot [101] (-B 1000).

### Identification and phylogenetic analysis of PKS genes in *Palythoa* species

We reconstructed metatranscriptome assemblies and MAGs using raw sequencing data. For metatranscriptome assembly, raw RNA-seq reads were treated with fastp as described above and assembled using rnaSPAdes v4.3.0 [102] with default settings. Amino acid sequences were obtained from resulting transcripts translated using TransDecoder with the option “-m 100”. For MAG reconstruction, raw HiFi reads were mapped to the host draft genomes, and unmapped HiFi reads were extracted. The extracted HiFi reads were assembled using metaMDBG v1.3.1 [103]. The metagenomic contigs were classified with Kraken2 v2.17.1 [104] against the standard database (v2026-06-26). Contigs assigned to Bacteria or Archaea (NCBI taxonomy IDs 2 and 2157) were extracted with KrakenTools v1.2.1 [105]. The contigs were binned with COMEBin v1.1.0 [106] using contig sequences and coverage. CheckM2 v1.1.0 [107] was used to estimate the quality of each bin. MAGs meeting at least the MIMAG medium-quality criteria (≥50% completeness and <10% contamination) [108] were retained. All retained MAGs were screened for biosynthetic gene clusters (BGCs) with antiSMASH v8.0.4 [109] with the option “--genefinding-tool prodigal”.

To screen PKS-like genes, we built a profile hidden Markov model (HMM) of the ketosynthase (KS) domain with hmmbuild of HMMER v3.4 [110] from the KS alignment [31]. We searched two sets of sequences with this HMM using hmmsearch (E-value cutoff: 1e-5). All hits at this E-value were treated as KS-domain sequences. (i) Genome assemblies of the five *Palythoa* species were translated in six frames with esl-translate (-l 100), so that genes missing from the gene annotations could also be found. (ii) The gene models and metatranscriptomes of each species were searched directly. For the metatranscriptome hits, we extracted the region of each protein aligned to the HMM. Redundant sequences, such as isoforms and the same transcript assembled in different runs, were removed with CD-HIT v4.8.1 [111] with options (“-c 0.99” and “-aS 0.9”).

We classified the candidate sequences with AFPK-Finder [31]. AFPK-Finder scores each sequence against 30 KS-related profile HMMs. A sequence is retained only if its full-sequence bit score exceeds 300 for at least one profile. The retained sequences are then embedded together with training KS sequences by t-SNE, at two perplexities.

For molecular phylogenetic analysis of PKS genes, as PKS/FAS are multidomain enzymes that are difficult to align, only ketosynthase (KS) domain (Beta-ketoacyl synthase-like, N-terminal domain, InterPro IPR014030; Beta-ketoacyl synthase, C-terminal domain, IPR014031) sequences were aligned as in Lin et al. [31], and molecular phylogenetic analyses were performed as described above. To reduce computational cost, we subsampled 30 genes from each clade using “sample” command of SeqKit v2.10.0 [112].

### Identification of symbiotic algal composition

The polyps from each colony were collected using clean scissors, rinsed in FSW, and placed in a 1.5-mL tube containing FSW. The polyp tissues were gently disrupted and squeezed against the tube wall with a pipette tip to release Symbiodiniaceae cells into FSW. The suspension was briefly centrifuged at low speed using a manual centrifuge (1011/1025, Hettich) to remove large tissue debris. The resulting supernatant containing Symbiodiniaceae cells was transferred to a new 1.5-mL tube and centrifuged (6000 × *g* for 5 min) to obtain Symbiodiniaceae cell pellets. Total DNA was extracted from the pellets using MightyPrep reagent for DNA (TaKaRa) according to the manufacturer’s instructions. The nuclear ITS2 region was amplified using Tks Gflex DNA Polymerase Low DNA (TaKaRa) with primers “SYM_VAR_5.8S” and “SYM_VAR_REV” [113]. The PCR conditions were as follows: an initial denaturation at 94 °C for 1 min, followed by 25 cycles of 98 °C for 10 s, 56 °C for 30 s, and 68 °C for 30 s, and a final extension step of 68 °C for 5 min. PCR products were pooled together in an equimolar concentration and sequenced on an Illumina MiSeq platform. Sequencing data were analyzed using the SymPortal pipeline [114] under default settings.

### Statistical analyses

Genome and gene statistics were calculated using SeqKit v2.10.0. Introns were defined as genomic regions between CDSs within genes. Pearson’s correlation coefficients were calculated using the “smplot2” package [115] and visualized with the “ggplot2” package [116] in R v4.5.1 [117].

For PGLS analyses, assembly size was regressed on each of the following predictors in separate models: TE length, non-TE length, class I length, class II length and unclassified repeat length. Models were fitted by maximum likelihood with “gls()” in the nlme v3.1-170 package [117]. The residual covariance structure followed Pagel’s λ model (“corPagel()”) [118, 119]. In this model, λ scales the phylogenetic covariances among species: λ = 0 corresponds to independent residuals (ordinary least squares) and λ = 1 to Brownian motion along the tree. The estimated λ was tested against λ = 0 and λ = 1 by likelihood-ratio tests (one degree of freedom), which assess, respectively, whether the residuals contain detectable phylogenetic signal and whether they depart from Brownian-motion expectations.

### Functional gene annotation

Functional annotation was performed using eggNOG-mapper v2.1.12 [120] based on the eggNOG v5.0 database [121]. Pcar was used as the representative zoantharian genome for clade III, and Pumb for clade II, based on genome BUSCO completeness. The actiniarian *Nematostella vectensis* was used as the anthozoan representative. Gene family enrichment analyses were conducted using topGO package [122], based on GO terms assigned by EggNOG-mapper. For survey of PKS genes, protein domain search using InterProScan [123] v5.69-97.0 was performed.

## Declarations

### Ethics approval and consent to participate

Not applicable

### Consent for publication

Not applicable

## Availability of data and materials

Raw sequencing data are deposited in the DRA under the accession numbers DRR932003–DRR932009 and DRR1213993-DRR1213994, and the genome assemblies of *P*. cf. *toxica* and *P*. sp. sakurajimensis are available in DDBJ/ENA/GenBank under accession numbers BAAKIW010000000 and BAAKIX010000000, respectively (BioProject PRJDB18008). The gene models for all species used in this study, metatranscriptome assemblies, and MAGs generated in this study are available in Zenodo (https://doi.org/10.5281/zenodo.23311753).

## Competing interests

The authors declare that they have no competing interests.

## Funding

This work was supported in part by grants from the Japan Society for the Promotion of Science (23H03822 to HY, 23H03821 to JDR).

## Authors’ Contributions

JDR and HY designed the research. YY, ES, CYL, MK, JDR and HY performed the research. YY analyzed the data and wrote the manuscript, with input from ES, JDR and HY. All authors reviewed and approved the final manuscript.

## Acknowledgements

We thank Dr. T. Naruse, T. van der Eeckhout, S. Borghi, B. Fiedler, and E. Conn (U. Ryukyus) for field support. We thank the Scientific Computing and Data Analysis Section in OIST for their computing resources. We would like to thank members of the Marine Genomics Unit at OIST. We thank all members of the Darwin Tree of Life Consortium for their commitment to making genome sequences openly accessible to the community.

## Notes

### Competing Interest Statement

The authors have declared no competing interest.

### Summary of Updates

Section on symbiosis and palytoxin updated with additional data.

