## Supplementary Materials for "Comparative genomics of zooxanthellate and azooxanthellate *Palythoa* (Cnidaria: Zoantharia) provides insights into symbiosis evolution and the origin of palytoxin"

Table S1. Summary of raw data generated in this study.

Table S2. Genome statistics of anthozoans used in this study.

Table S3. Gene model statistics of anthozoans used in this study.

Table S4. Summary of orthogroup identification.

Table S5. List of orthogroups and the number of genes in each species.

Table S6. Orthogroups significantly expanded in the common ancestor of *Palythoa*.

Table S7. Orthogroups restricted to *Palythoa*.

Table S8. Orthogroups putatively lost in *Palythoa*.

Table S9. Orthogroups specific to each *Palythoa* clade.

Table S10. Fast-evolving genes in each *Palythoa* clade.

Table S11. The output of SymPortal.

Table S12. Summary of metagenome assemblies constructed from HiFi reads of *Palythoa* species.

Table S13. Quality and taxonomic classification of medium-quality metagenome-assembled genomes (MAGs) recovered from *Palythoa* species.

Table S14. Summary of metatranscriptome assemblies in *Palythoa*.

Table S15. Repeat content of the 16 anthozoan genome assemblies used in the PGLS analysis.

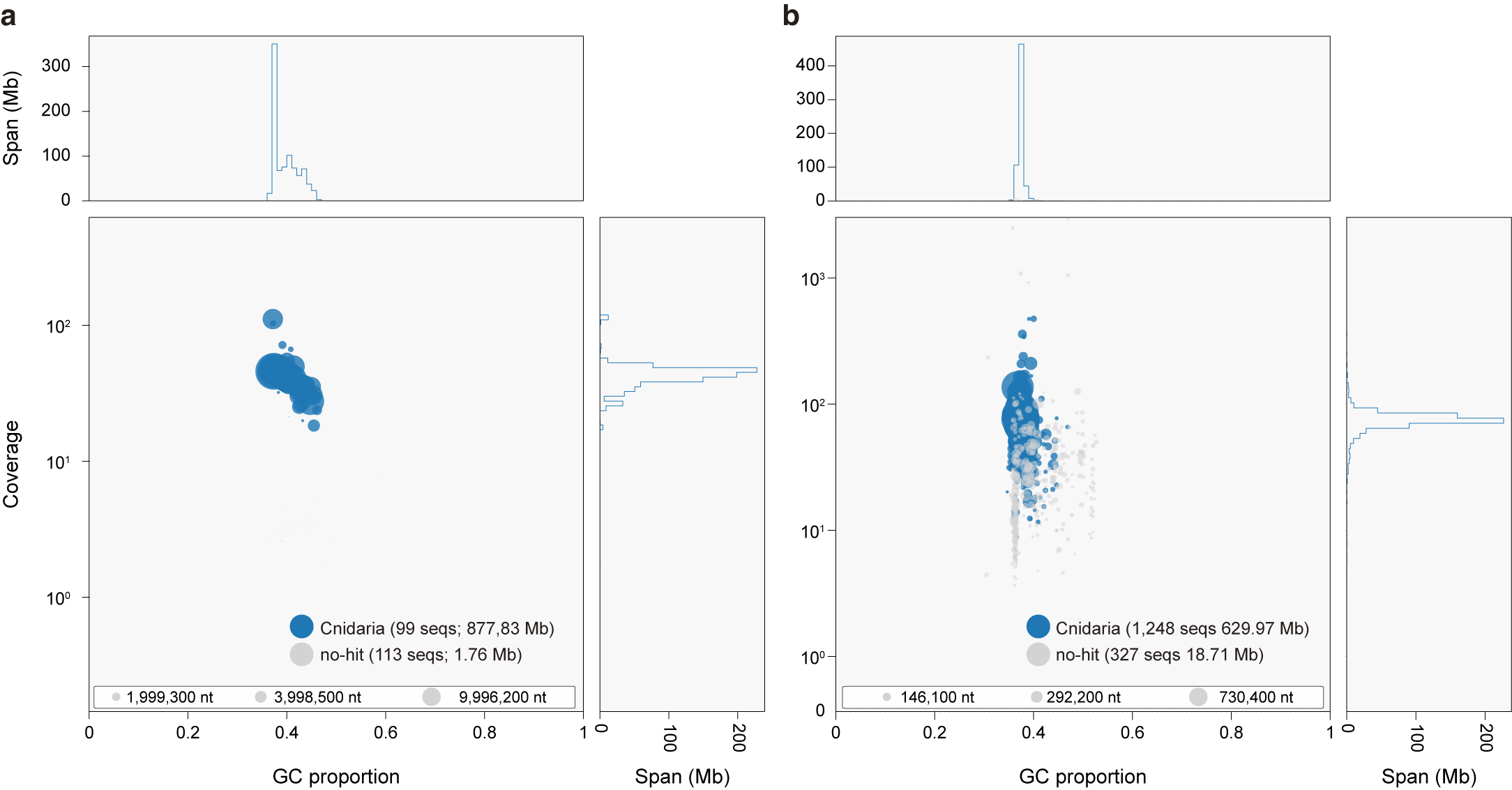

**Fig. S1. BlobTools plots of the genome assemblies of (a) *Palythoa* sp. sakurajimensis and (b) *P.* cf. *toxica*.**

Contigs are plotted by GC content and read coverage, with circle size proportional to contig length and colors indicating taxonomic assignment. All contigs were assigned to Cnidaria or had no hit, indicating no detectable contamination.

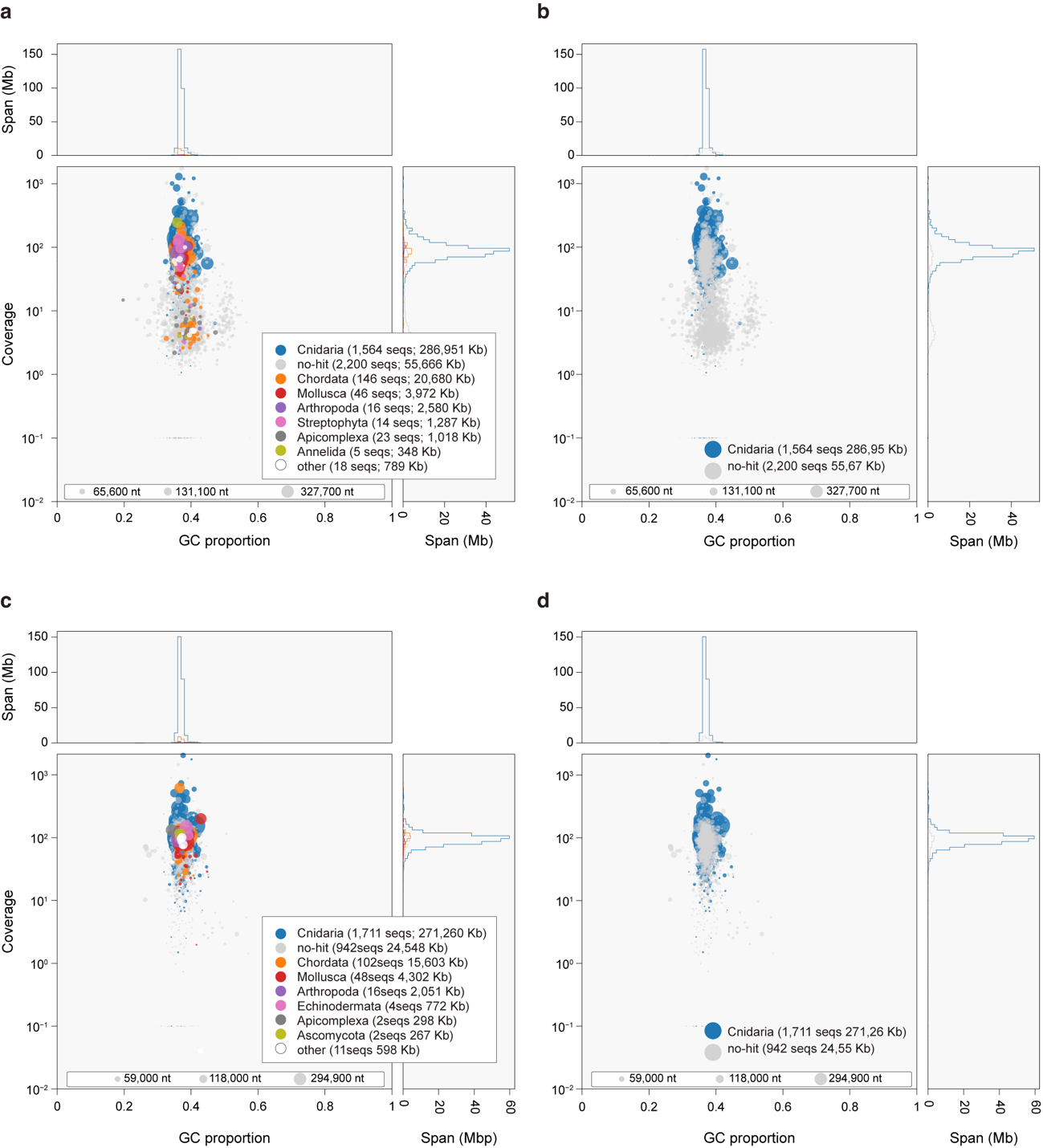

**Fig. S2. BlobTools plots of the *Palythoa* *mizigama* and *P. umbrosa* assemblies before and after removal of contaminant contigs.**

(**a, b**) *P. mizigama* before (**a**) and after (**b**) cleaning. (**c, d**) *P. umbrosa* before (**c**) and after (**d**) cleaning. Contigs assigned to taxa other than Cnidaria were removed. Plot elements are as in Fig. S1.

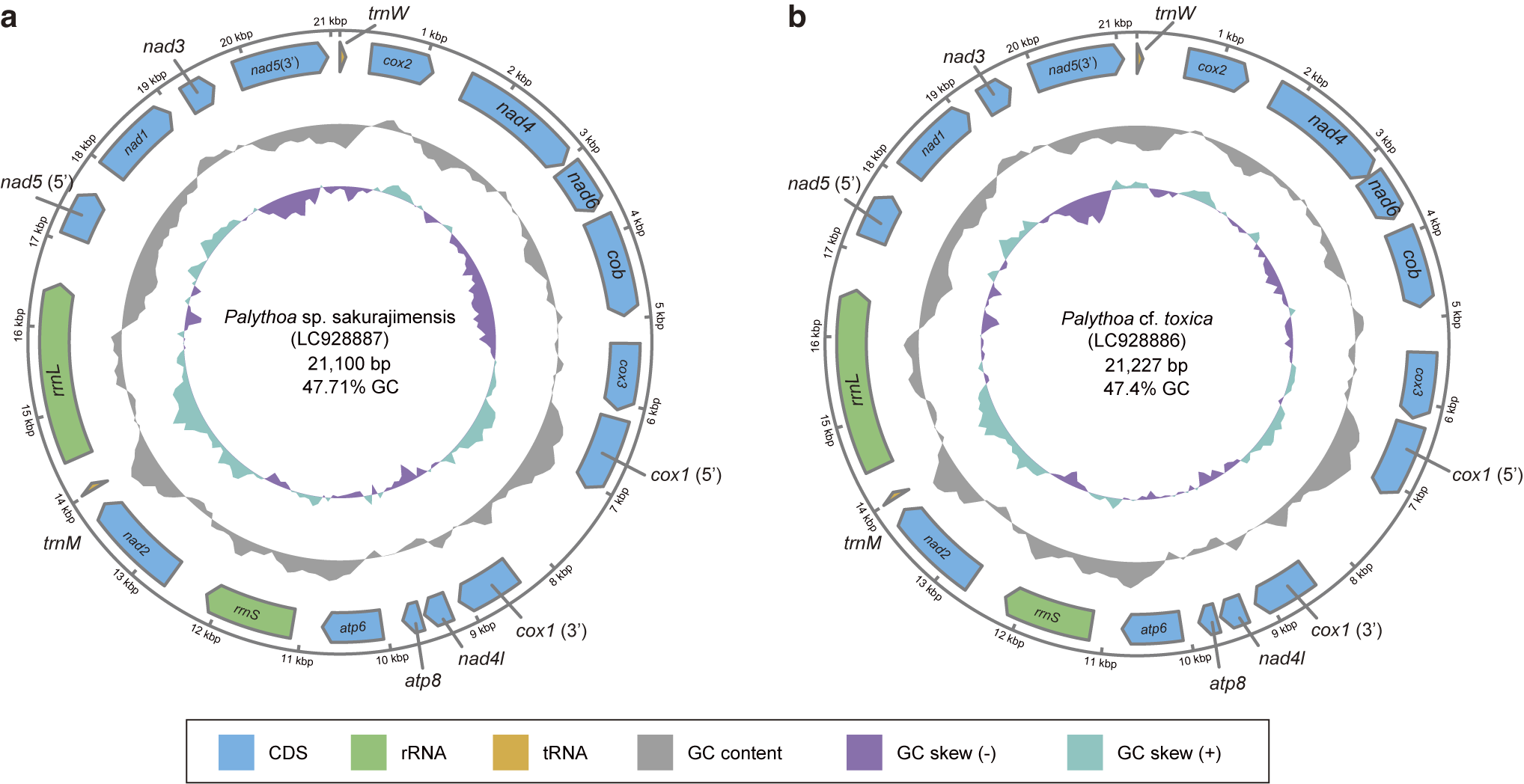

**Fig. S3. Complete mitochondrial genomes of (a) *Palythoa* sp. sakurajimensis and (b) *P.* cf. *toxica*.**

Genes are colored by type (protein-coding, rRNA, and tRNA genes), and the inner rings show GC content and GC skew. Accession numbers, genome sizes, and GC contents are shown at the center.

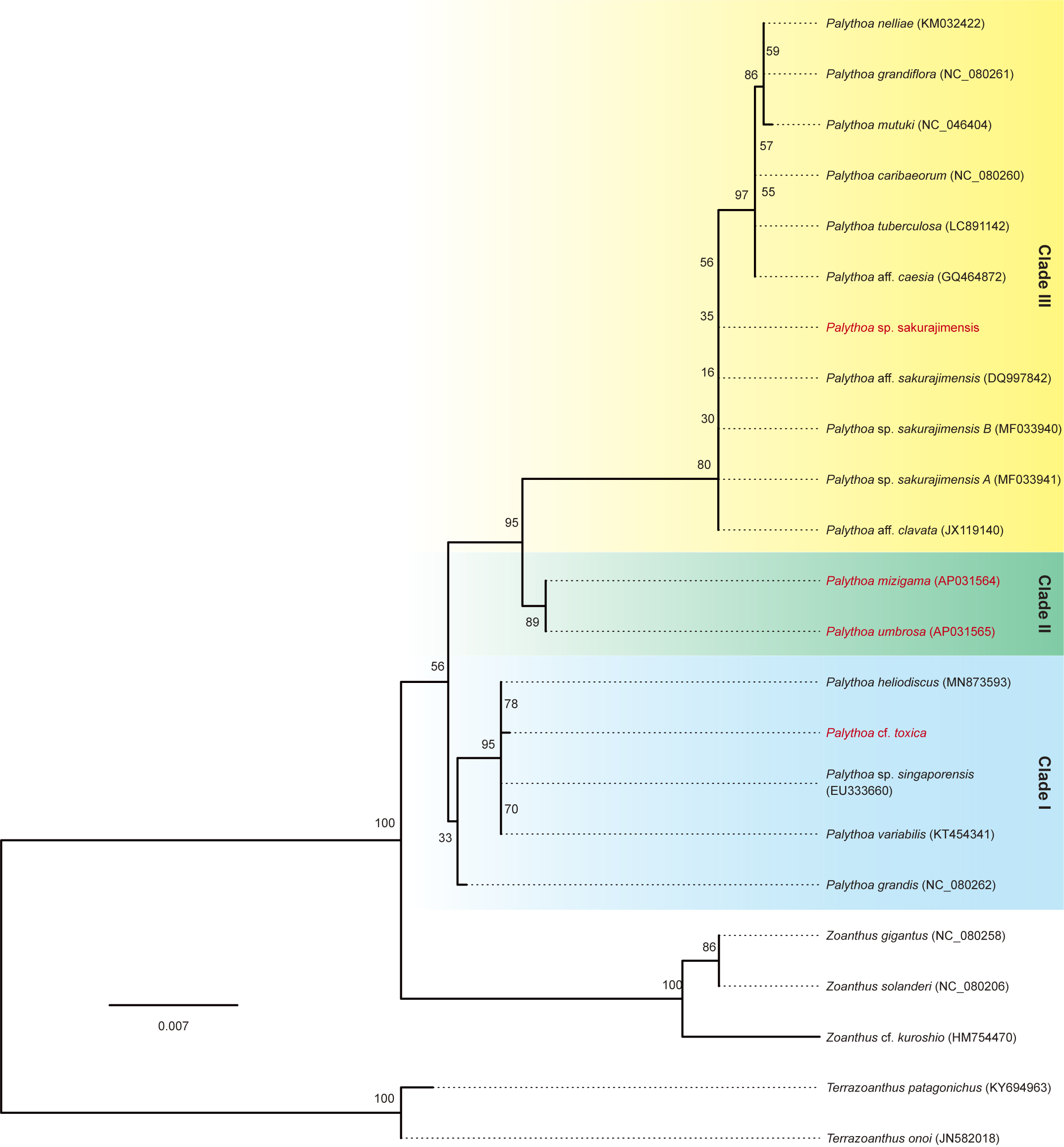

**Fig. S4. Maximum-likelihood phylogeny of *Palythoa* based on partial mitochondrial *rrnL* sequences.**

The tree was inferred from 2,093 nucleotide sites with IQ-TREE v2.2.2.6 under the best-fit model TN+F+I. Species analyzed in this study are shown in red, and GenBank accession numbers are given in parentheses. Clades I–III correspond to those in the main text. *Zoanthus* and *Terrazoanthus* sequences were used as outgroups. Numbers at nodes indicate bootstrap values (1,000 replicates), and the scale bar indicates substitutions per site.

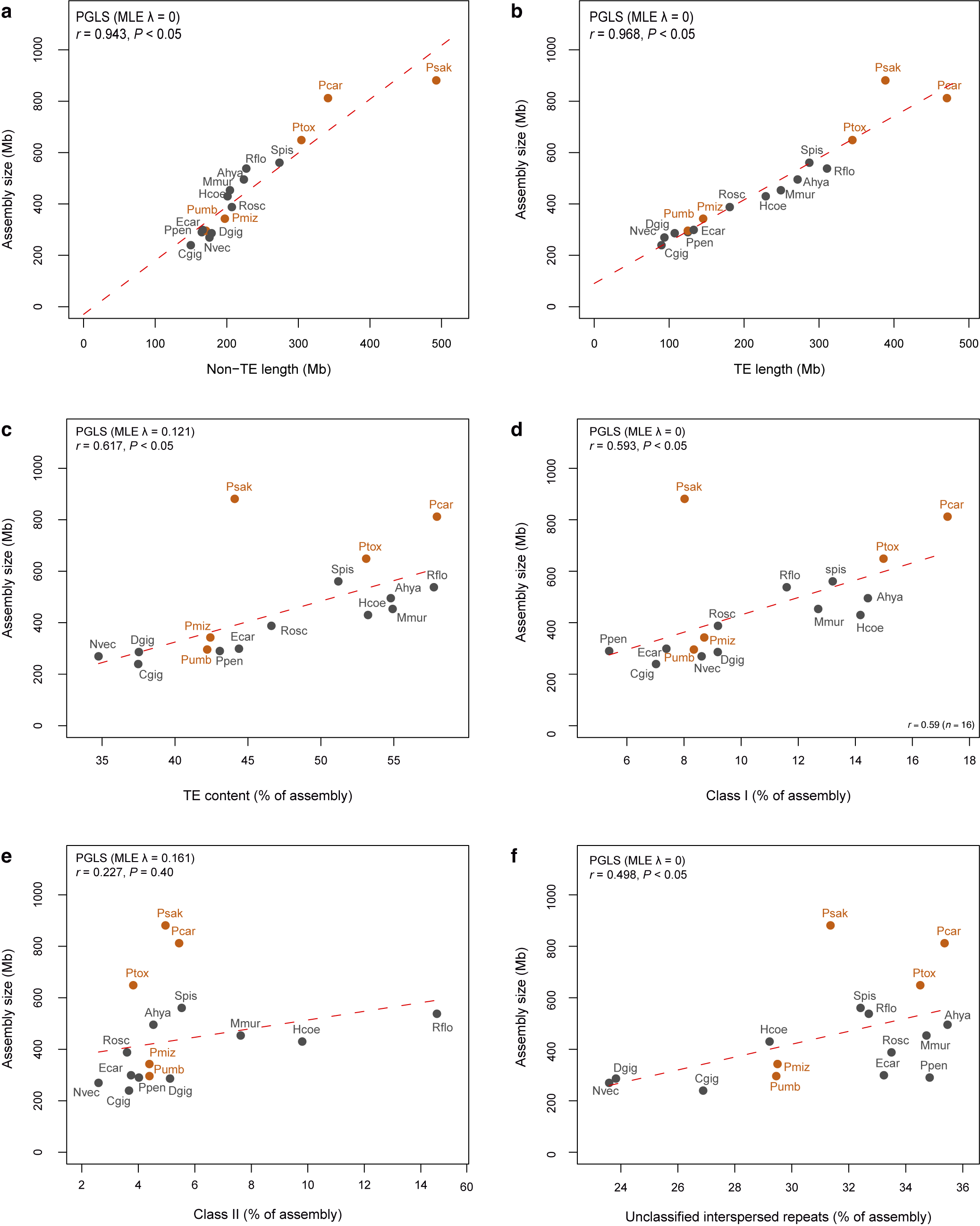

**Fig. S5. Relationships between assembly size and repeat content in anthozoans.**

Assembly size is plotted against (**a**) non-TE length (assembly size minus the total length of interspersed repeats) and (**b**) TE length (total length of interspersed repeats), and the proportion of each assembly occupied by (**c**) all TEs, (**d**) class I elements (LINEs, SINEs, LTRs, PLEs, and retroposons), (**e**) class II elements (DNA transposons and rolling-circle/Helitron elements), (**f**) unclassified interspersed repeats, for 16 anthozoan genome assemblies. Orange and grey circles indicate *Palythoa* and other anthozoan species, respectively; species abbreviations are listed in Table S2. Red dashed lines indicate regressions fitted by phylogenetic generalized least squares (PGLS) on the ultrametric species tree derived from Fig. 1d, with Pagel's λ estimated by maximum likelihood (shown in each panel). *r* and *P* in each panel indicate Pearson's correlation coefficients and their significance. Values for all species are listed in Table S15.

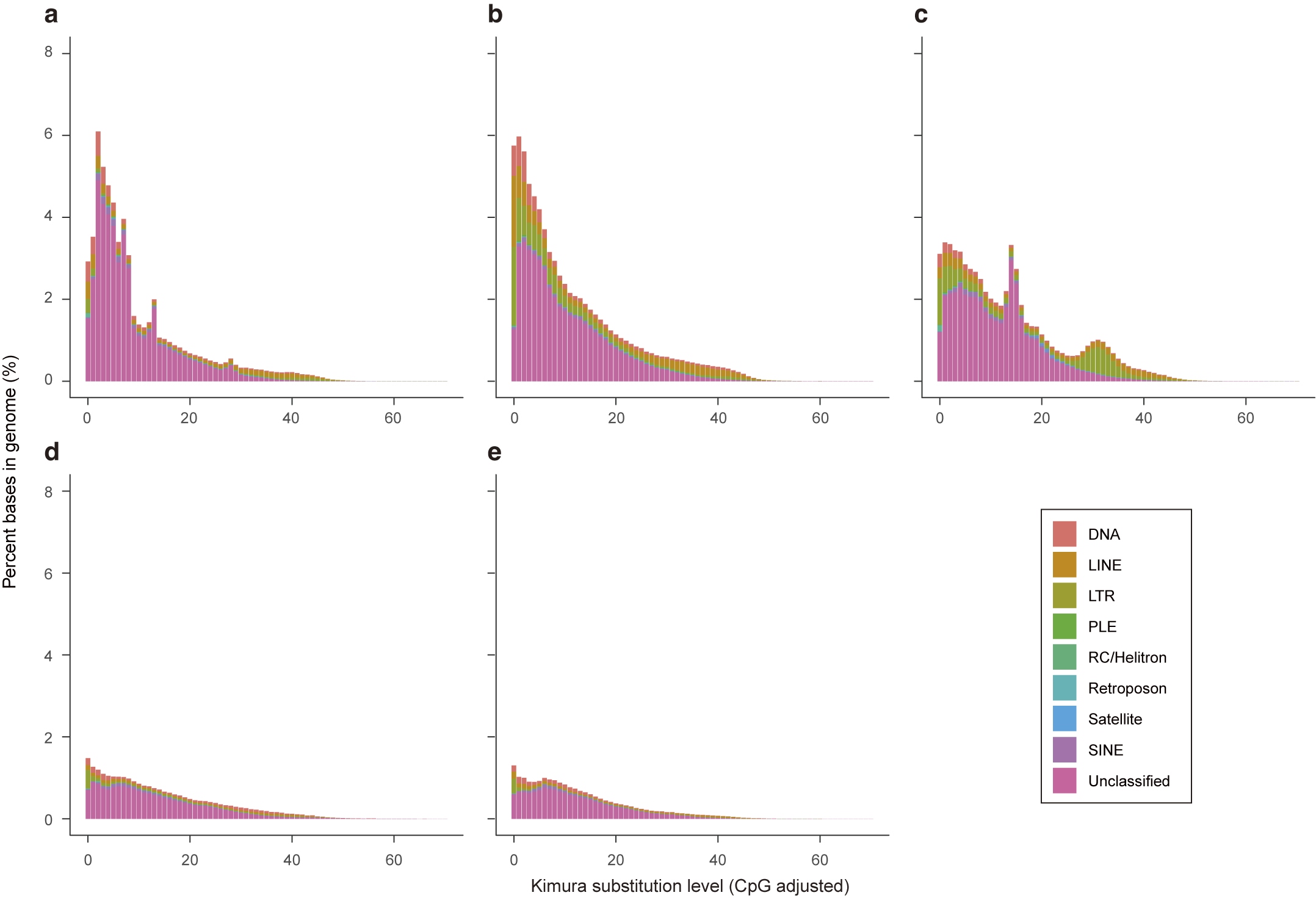

**Fig. S6. Repeat landscapes of *Palythoa* genomes.**

(**a**) *P.* sp. sakurajimensis, (**b**) *P. caribaeorum*, (**c**) *P.* cf. toxica, (**d**) *P. mizigama*, and (**e**) *P. umbrosa*. Bars show the percentage of each genome occupied by TE copies at each level of Kimura divergence (CpG-adjusted) from their consensus sequences, colored by TE category. Copies with low divergence likely represent recent insertions, whereas those with high divergence represent older, degenerated copies.

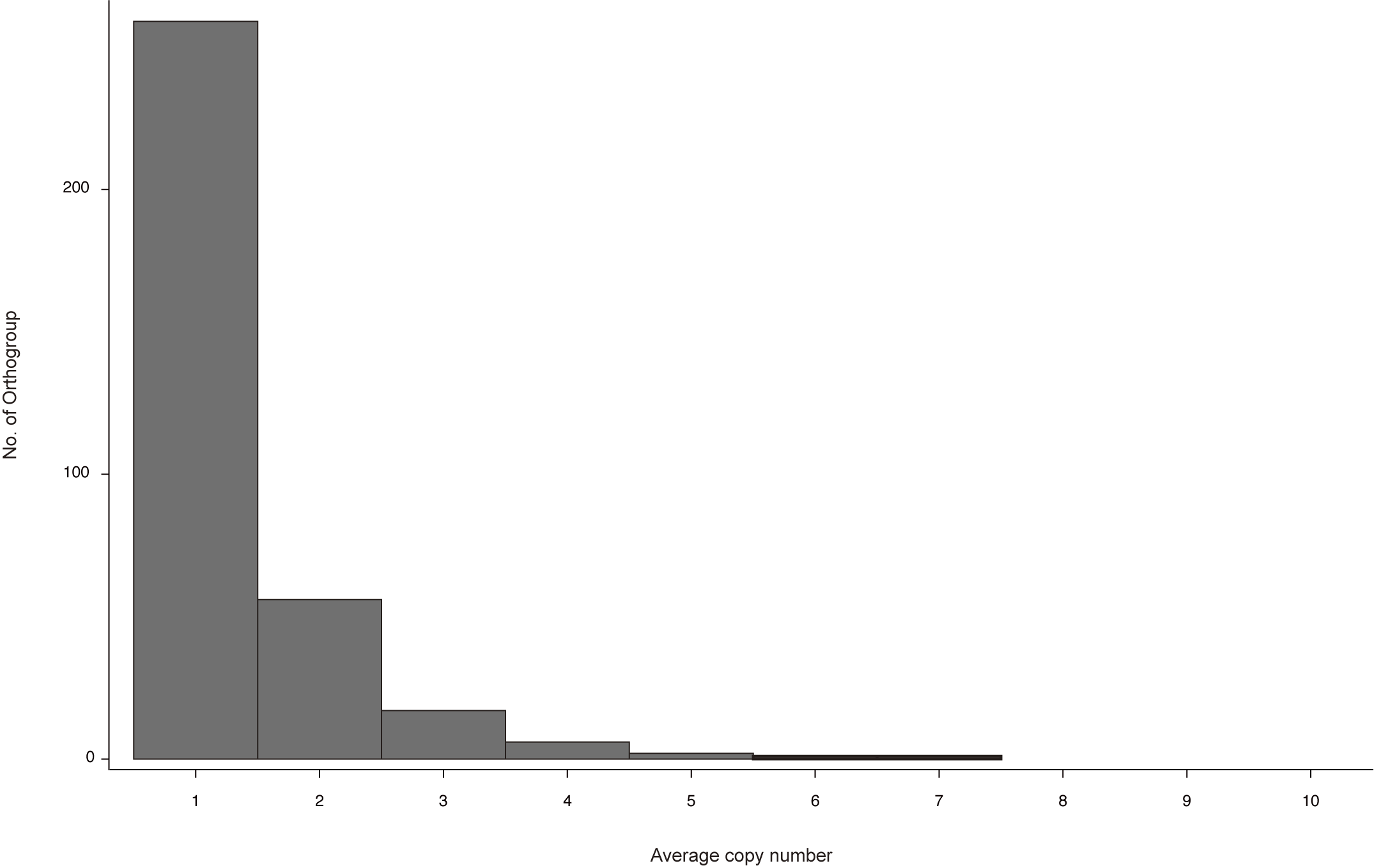

**Fig. S7. Mean copy numbers of *Palythoa*-restricted orthogroups (OGs).**

Distribution of the mean number of gene copies per species for the 342 OGs restricted to zoantharians.

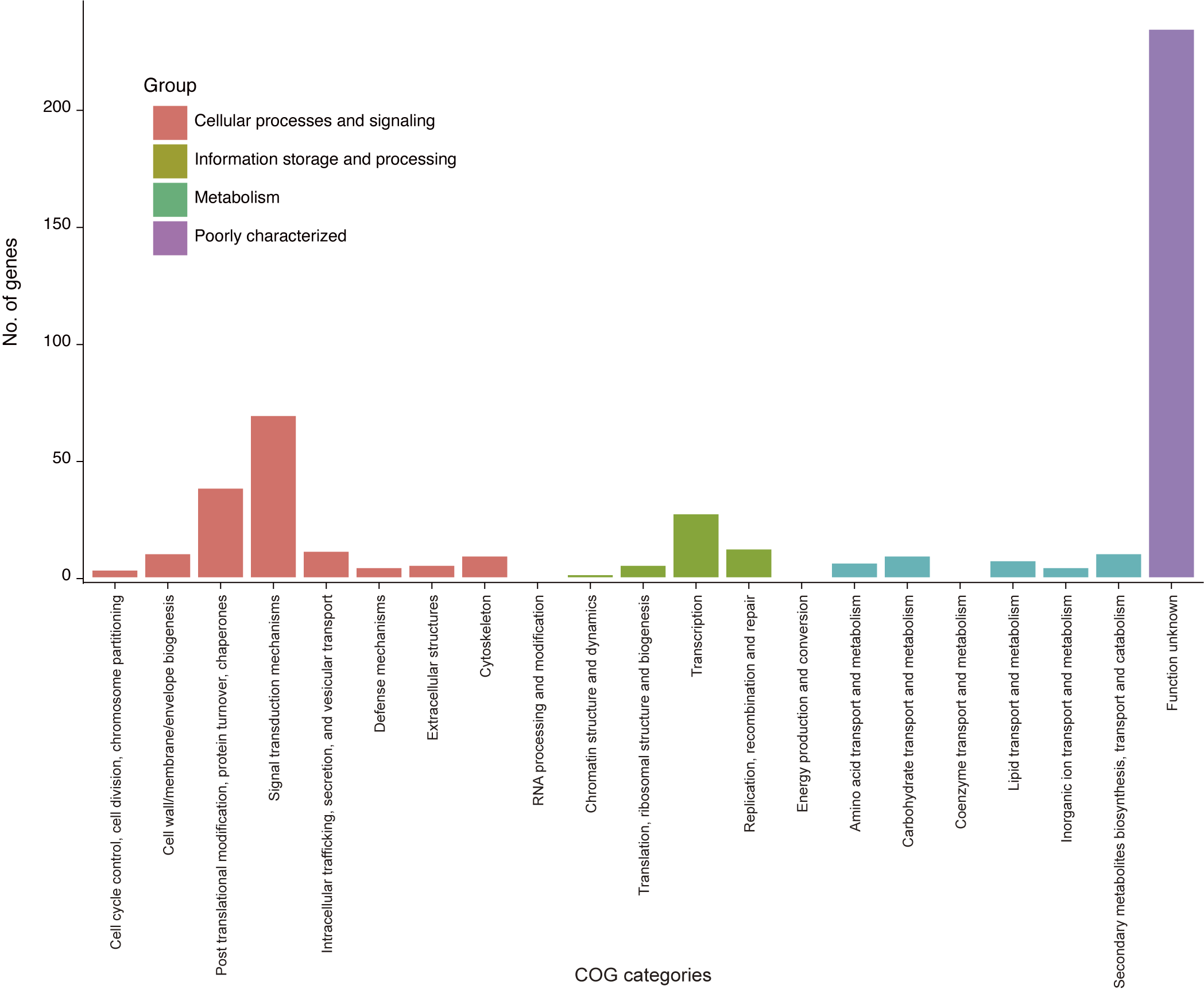

**Fig. S8. Functional categories of genes in *Palythoa*-restricted orthogroups.**

COG categories were assigned to *Palythoa caribaeorum* genes, used as representatives, with eggNOG-mapper v2.1.12 based on the eggNOG v5.0 database. Colors indicate COG functional groups.

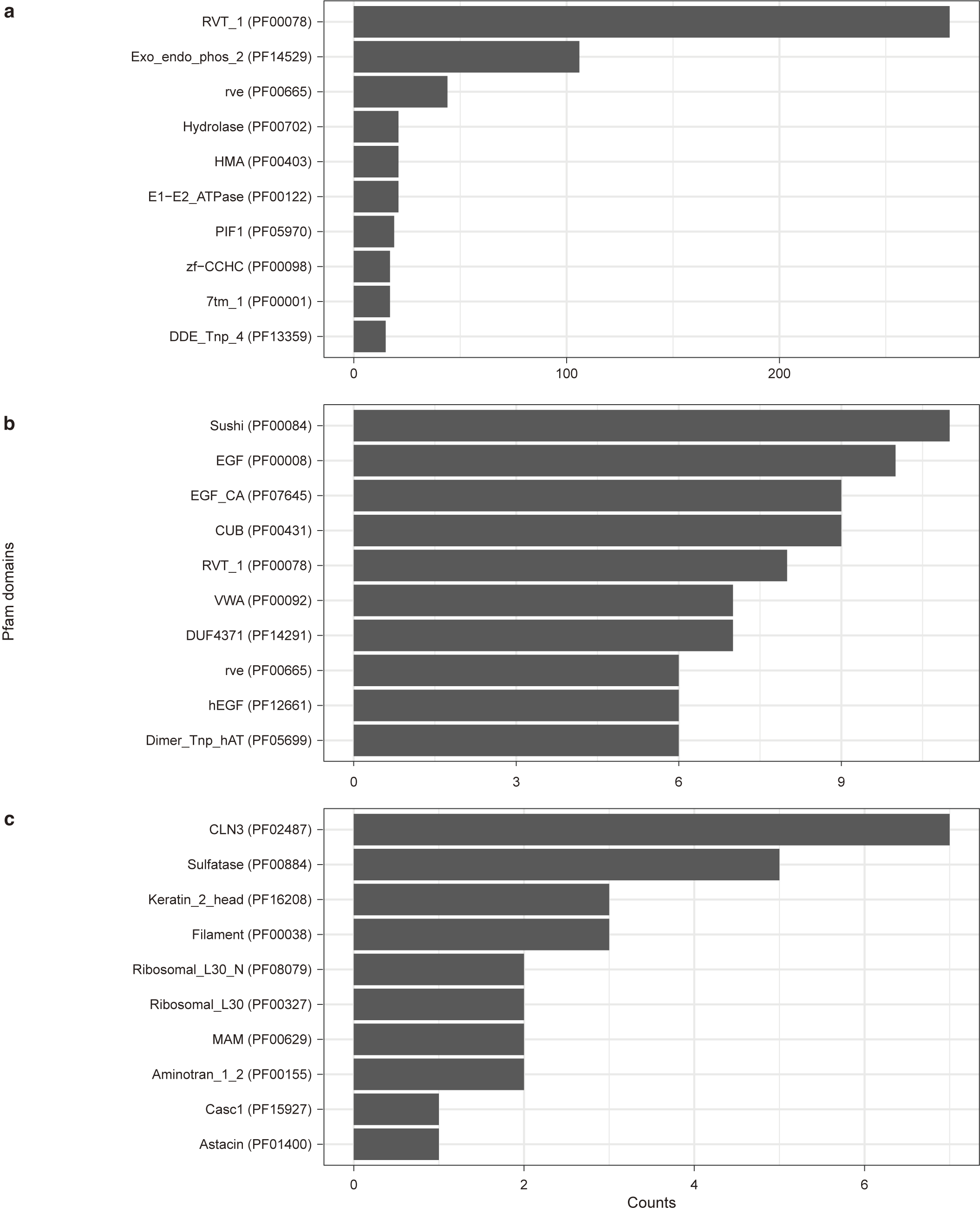

**Fig. S9. The ten most frequent Pfam domains in clade-specific orthogroups of *Palythoa*.**

(**a**) Clade I (*P.* cf. *toxica*), (**b**) clade II (*P. umbrosa*), and (**c**) clade III (*P. caribaeorum*). Counts indicate the number of genes containing each domain. Pfam domains were assigned with eggNOG-mapper v2.1.12 based on the eggNOG v5.0 database.

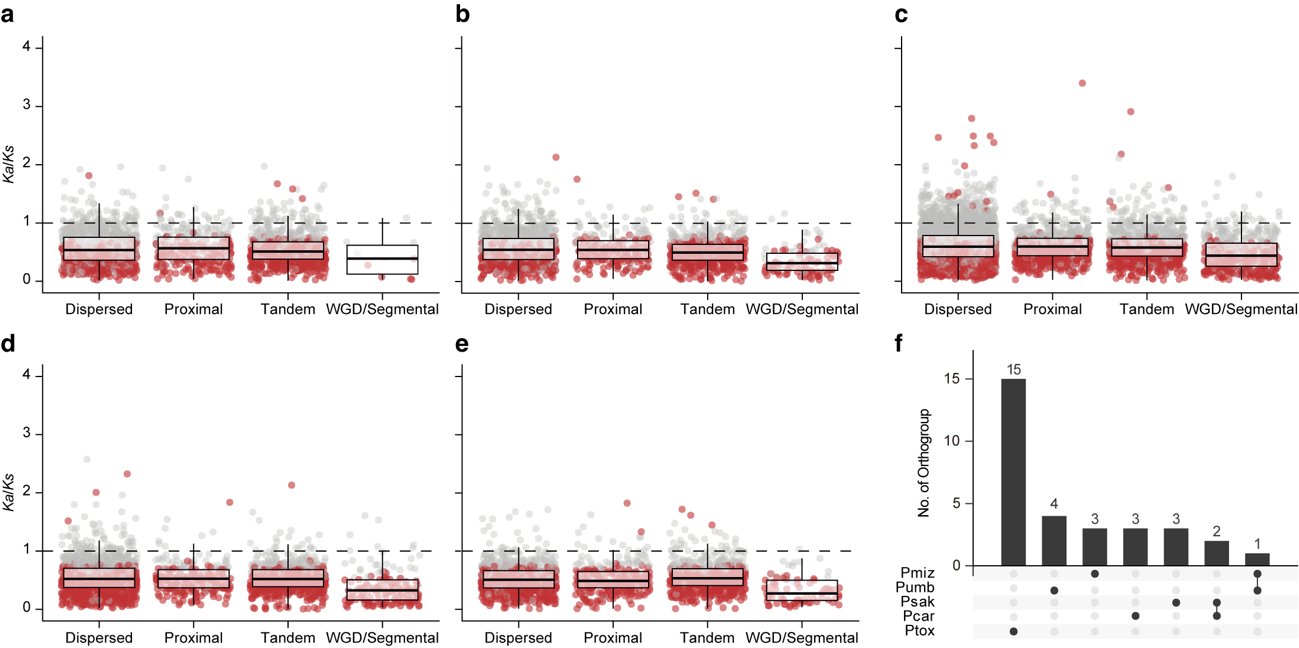

**Fig. S10. Paralogous gene pairs under positive selection in *Palythoa*.**

(**a–e**) *Ka*/*Ks* ratios of paralogous gene pairs in each duplication mode in (**a**) *P. caribaeorum*, (**b**) *P.* sp. sakurajimensis, (**c**) *P.* cf. *toxica*, (**d**) *P. mizigama*, and (**e**) *P. umbrosa*. Boxes indicate medians and interquartile ranges. Red points indicate pairs whose *Ka*/*Ks* ratios differ significantly from 1 (Benjamini–Hochberg-adjusted *P* < 0.05), and pairs above the dashed line (*Ka*/*Ks* = 1) were considered to be under positive selection. (**f**) UpSet plot showing the numbers of orthogroups containing positively selected gene pairs in each species and shared among species.

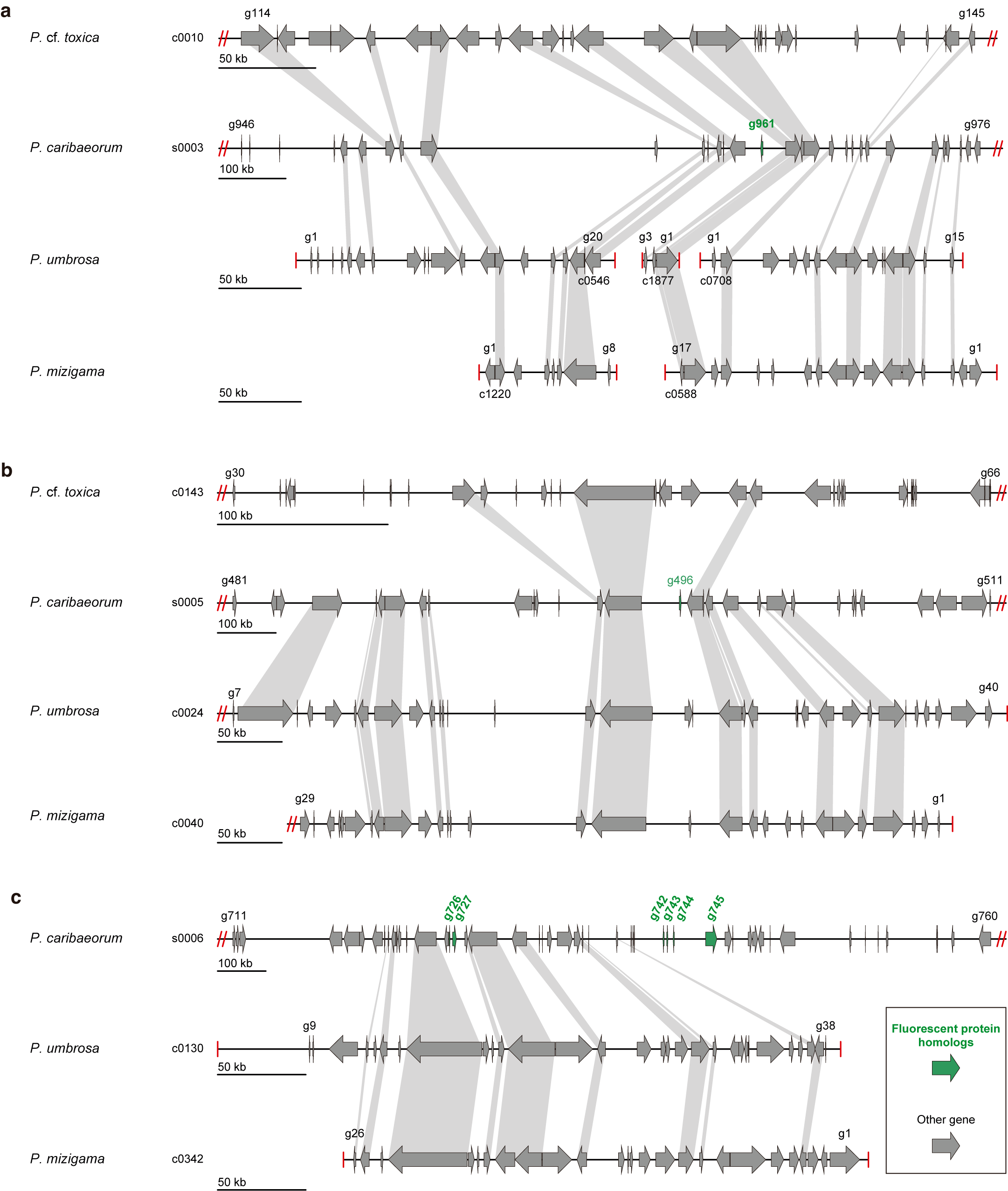

**Fig. S11. Microsynteny of genomic regions surrounding GFP-like fluorescent protein genes in Palythoa.**

Genomic regions containing genes encoding GFP-like fluorescent proteins (OG0000497; green arrows) in the chromosome-level assembly of the zooxanthellate Palythoa caribaeorum were used as references and compared with the corresponding regions of the azooxanthellate P. umbrosa and P. mizigama. (**a**) Region surrounding g961 on scaffold s0003, together with the corresponding region of the zooxanthellate P. cf. toxica (contig c0010). (**b**) Region surrounding g496 on scaffold s0005, together with the corresponding region of the zooxanthellate P. cf. toxica (contig c0143). (**c**) Region surrounding g726, g727 and g742–g745 on scaffold s0006. Arrows represent predicted protein-coding genes and their transcriptional orientation; GFP-like genes are shown in green and all other genes in grey. Grey bands connect homologous genes, defined as genes assigned to the same orthogroup. Labels at the ends of each track give the first and last genes displayed; genes are numbered sequentially along each contig or scaffold, and tracks in which numbers decrease from left to right are shown in reverse-complement orientation. Red double slashes indicate that the sequence continues beyond the displayed region, and red vertical bars indicate contig ends. Each track has its own scale bar. In (**b**) and around g726–g727 in (**c**), genes flanking the GFP-like genes are retained on single contigs in both azooxanthellate species, but no GFP-like gene is present between them and no homologous sequence was detected by BLASTN and HMM searches of six-frame translations of each genome assembly. In (**a**) and around g742–g745 in (**c**), the corresponding regions are interrupted by contig ends in the azooxanthellate assemblies, so the presence or absence of GFP-like genes cannot be determined from synteny alone.

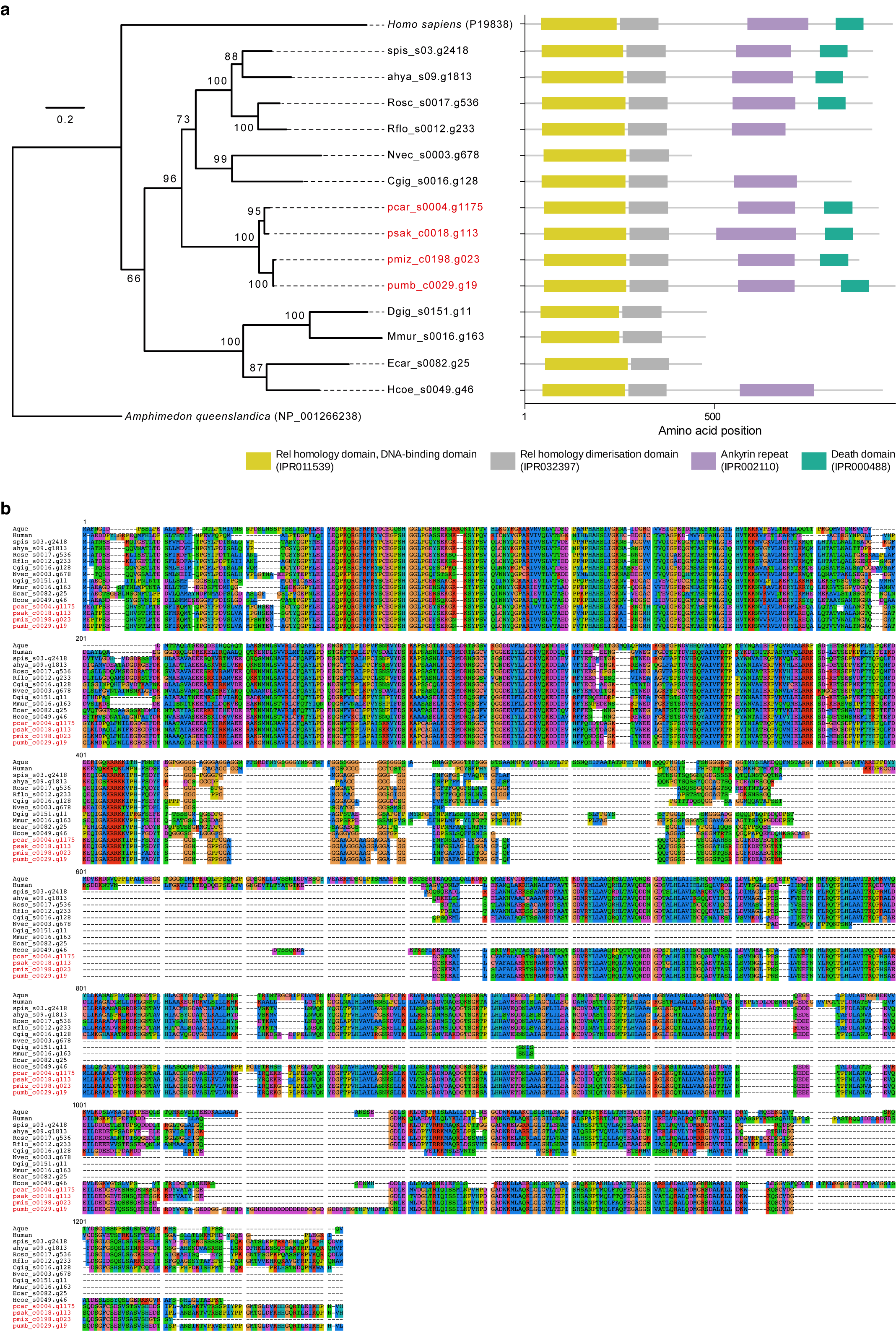

**Fig. S12. Phylogeny and domain architecture of NF-κB homologs in anthozoans.**

(**a**) Maximum-likelihood phylogeny (left) and domain architecture (right) of NF-κB (Rel family) proteins. *Palythoa* sequences are shown in red. The tree was inferred from 1,077 amino acid sites with IQ-TREE v2.2.2.6 under the best-fit model Q.pfam+F+I+G4 and rooted with NF-κB of the sponge *Amphimedon queenslandica* (NP_001266238), and human NF-κB1 (P19838) is included as a reference. Numbers at nodes are ultrafast bootstrap support values (1,000 replicates). Scale bar, 0.2 substitutions per site. Domains were annotated with InterProScan: Rel homology DNA-binding domain (IPR011539), Rel homology dimerization domain (IPR032397), ankyrin repeats (IPR002110) and death domain (IPR000488). (**b**) Multiple sequence alignment of the NF-κB proteins shown in (**a**), generated with MAFFT v7.525 and visualized with SeaView. Numbers above the blocks indicate alignment positions. All four Palythoa species examined, including the azooxanthellate P. mizigama and P. umbrosa, retain the complete domain architecture, and no amino acid substitutions specific to the azooxanthellate species were detected within conserved domains. No NF-κB homolog was detected in P. cf. toxica by BLASTN or HMM searches of six-frame translations of its genome assembly; given the lower contiguity and completeness of this assembly, we cannot distinguish between gene loss and incomplete assembly. Gene identifiers consist of a species abbreviation, contig or scaffold, and gene number; species abbreviations are listed in Table S2.

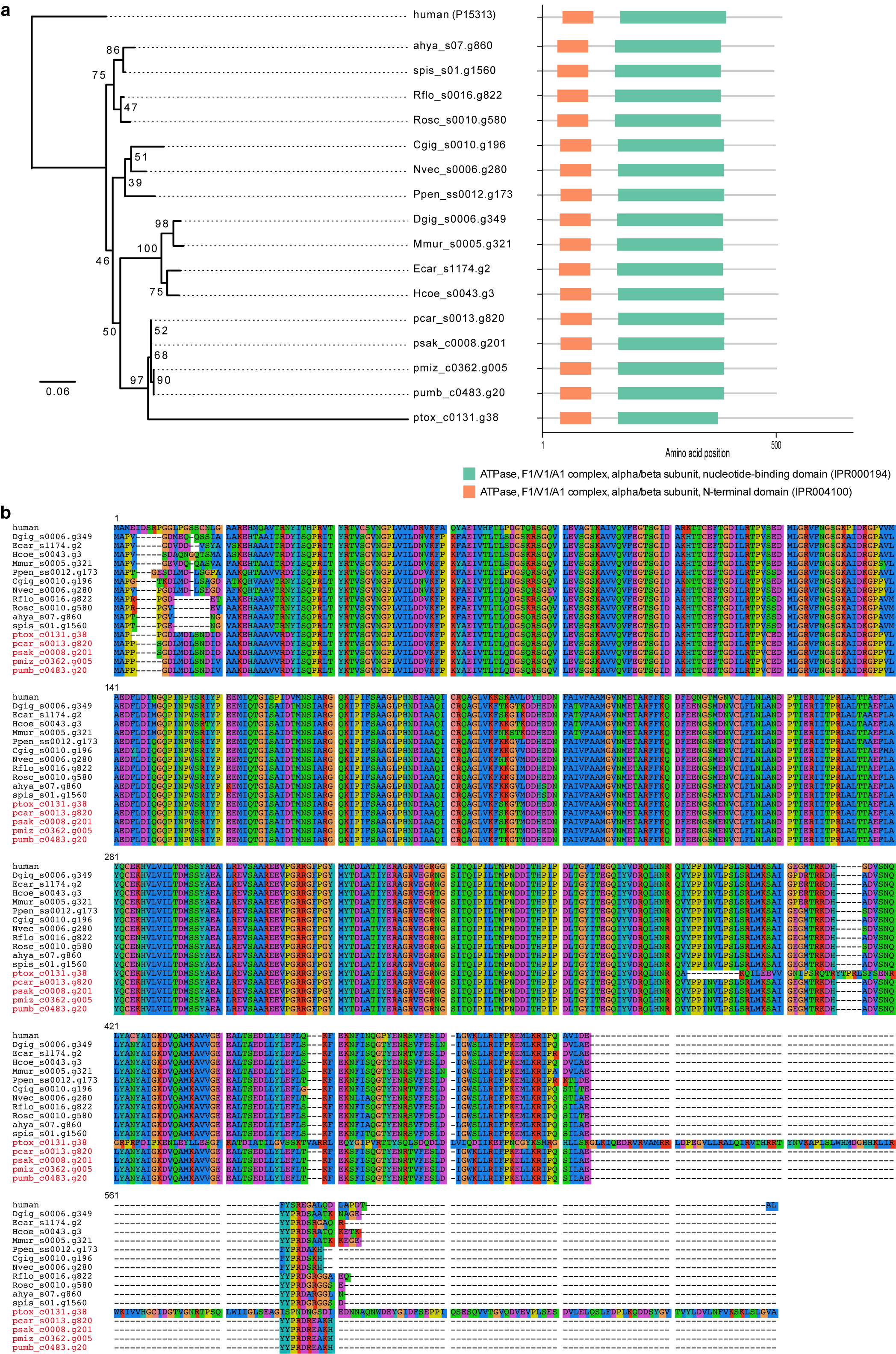

**Fig. S13. Phylogeny, domain architecture, and sequence alignment of V-type H⁺-ATPase subunit B (VHAB) homologs in anthozoans.**
(**a**) Maximum-likelihood phylogeny (left) and domain architecture (right) of VHAB proteins. The tree was inferred from an alignment of 510 amino acid sites with IQ-TREE v2.2.2.6 under the VT+I+R3 model selected by ModelFinder according to BIC, and rooted with human VHAB (P15313). Palythoa sequences are shown in red. Numbers at nodes are ultrafast bootstrap support values (1,000 replicates). Scale bar, 0.06 substitutions per site. Domains were annotated with InterProScan: ATPase, F1/V1/A1 complex, alpha/beta subunit, N-terminal domain (IPR004100) and nucleotide-binding domain (IPR000194). (**b**) Multiple sequence alignment of the VHAB proteins shown in (**a**), generated with MAFFT v7.525 and visualized with SeaView. Numbers above the blocks indicate alignment positions. VHAB was identified in all five Palythoa species, all of which retain both domains. No azooxanthellate-specific substitutions were found at conserved sites. Gene identifiers consist of a species abbreviation, contig or scaffold, and gene number; species abbreviations are listed in Table S2.

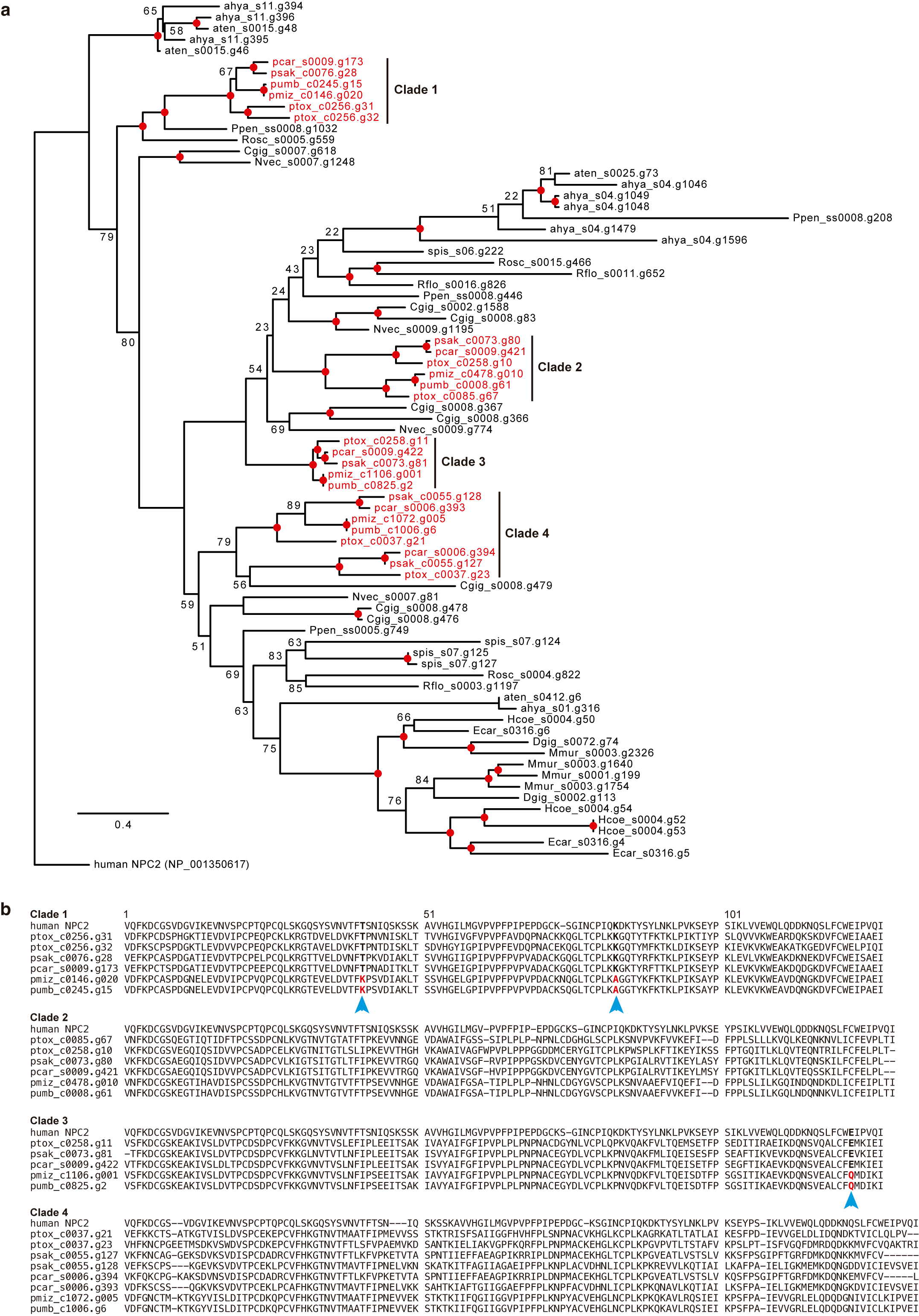

**Fig. S14. Phylogeny and sequence comparison of NPC2 homologs in *Palythoa*.**

(**a**) Maximum-likelihood phylogeny of Niemann–Pick type C2 (NPC2) proteins from anthozoans. The tree was inferred from an alignment of 152 amino acid sites with IQ-TREE v2.2.2.6 under the WAG+F+I+G4 model selected by ModelFinder according to BIC, and rooted with human NPC2 (NP_001350617). *Palythoa* sequences (red) fall into four clades (Clades 1–4). Red circles indicate nodes with ultrafast bootstrap support ≥ 90% (1,000 replicates); support values for other nodes are shown as numbers. Scale bar, 0.4 substitutions per site. Gene identifiers consist of a species abbreviation, contig or scaffold, and gene number; species abbreviations are listed in Table S2. (**b**) Amino acid alignments of *Palythoa* NPC2 proteins in each clade with human NPC2. Numbers above the alignment indicate alignment positions. Blue arrowheads and red letters mark positions where the azooxanthellate species (*P. mizigama* and *P. umbrosa*) carry residues different from those of all zooxanthellate *Palythoa* species (two positions in Clade 1 and one in Clade 3); residues of the other sequences at these positions are shown in bold. No such positions were found in Clades 2 and 4.

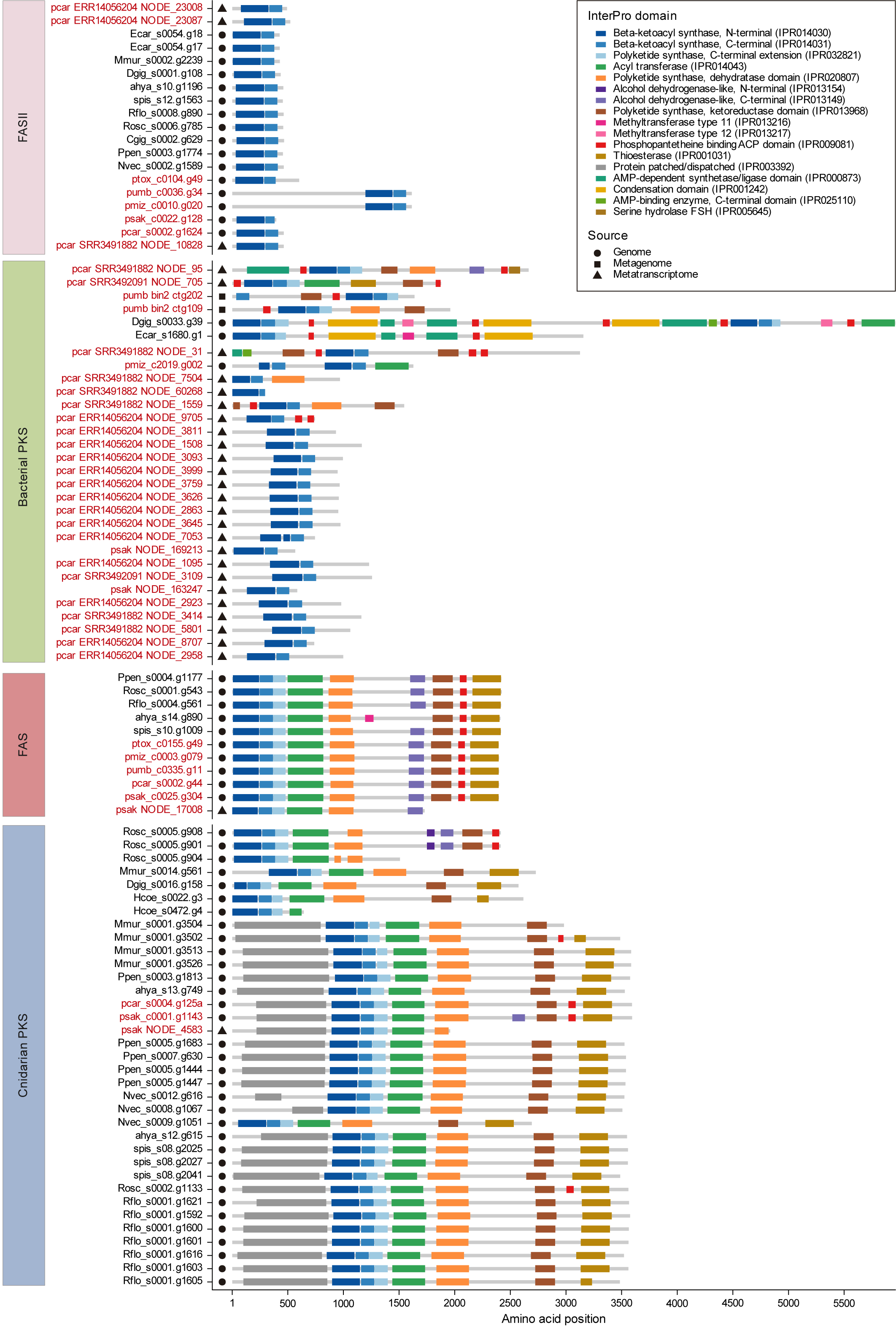

**Fig. S15. Domain architectures of ketosynthase domain-containing proteins from cnidarian genomes, metagenomes, and metatranscriptomes.**

Proteins containing ketosynthase (KS) domains were classified into four groups: type II fatty acid synthase (FASII), bacterial-like polyketide synthase (PKS), type I fatty acid synthase (FAS), and cnidarian PKS, based on the KS domain phylogeny (Fig. 6). Gray bars represent full-length proteins, and colored boxes indicate protein domains annotated by using InterProScan. The x-axis shows amino acid positions. Symbols indicate the data source: circles, gene models predicted from genome assemblies; squares, genes from metagenome-assembled bins; triangles, genes from metatranscriptome assemblies. Sequence IDs shown in red are derived from *Palythoa* species (pcar, *P. caribaeorum*; pmiz, *P. mizigama*; ptox, *P.* cf. *toxica*; psak, *Palythoa* sp. sakurajimensis; pumb, *P. umbrosa*). For metagenome- and metatranscriptome-derived sequences, IDs consist of the sequencing run accession or bin name followed by the contig ID. Species abbreviations are listed in Table S2.

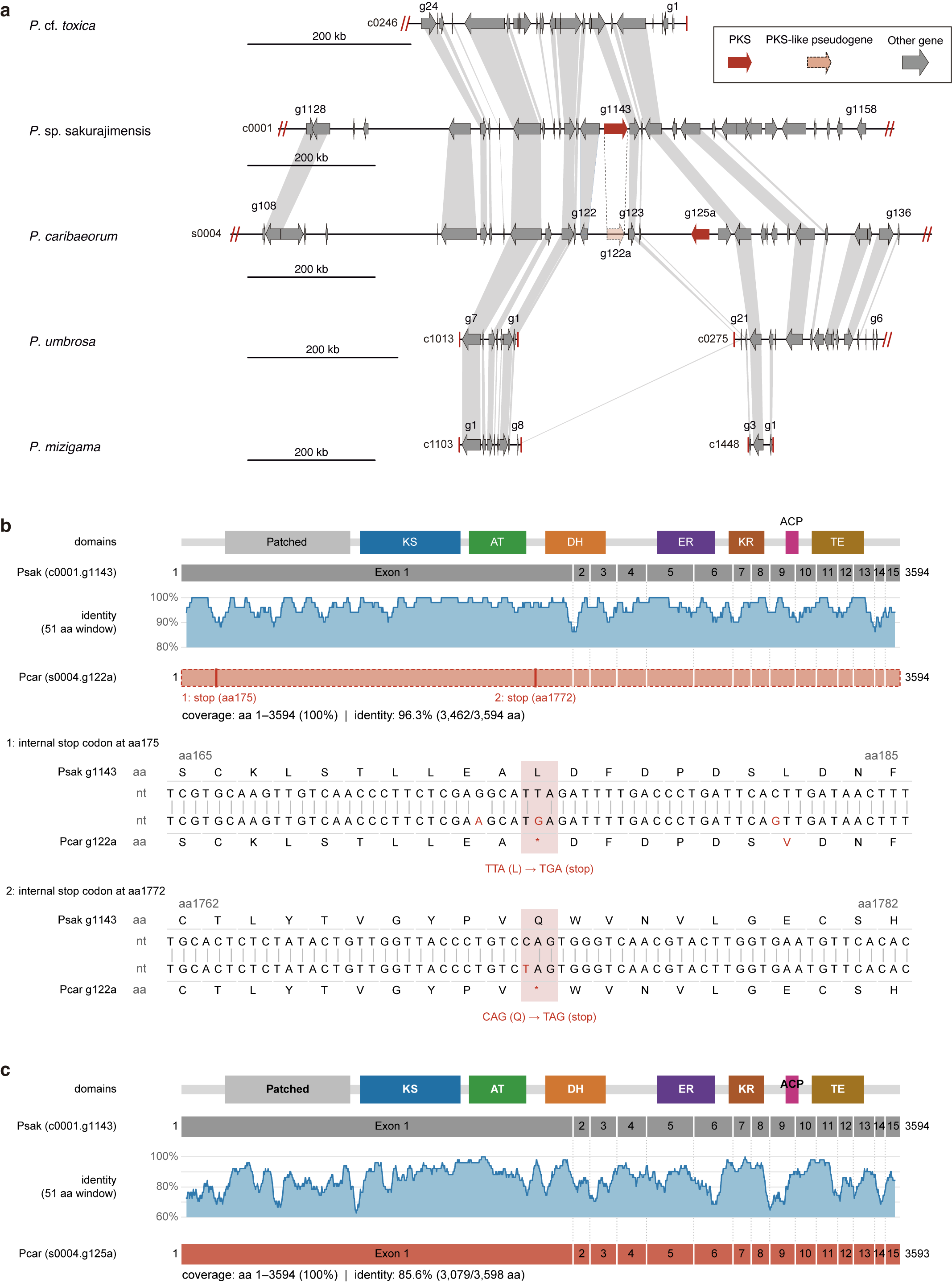

**Fig. S16. Microsynteny and sequence comparison of polyketide synthase (PKS) genes in *Palythoa*.**

(**a**) Microsynteny of the genomic region surrounding the PKS gene of *P.* sp. sakurajimensis (c0001.g1143) and the corresponding regions in *P.* cf. *toxica*, *P. caribaeorum*, *P. umbrosa* and *P. mizigama*. Red arrows indicate PKS genes, the dashed salmon arrow indicates a PKS pseudogene, and grey arrows indicate other genes; arrow direction shows transcriptional orientation. Grey bands connect homologous genes, defined as genes assigned to the same orthogroup, and dotted lines link g1143 to its syntenic counterpart in *P. caribaeorum* (s0004.g122a). Gene identifiers with the suffix “a” (g122a and g125a) denote loci that were absent from the original gene models and were identified by a genome-wide search of six-frame translations. Of these, g125a was added to the gene models, whereas g122a was not, because it contains two premature stop codons and is likely a pseudogene (see b). Labels give the first and last genes displayed; tracks in which gene numbers decrease from left to right are shown in reverse-complement orientation. Red double slashes indicate that the sequence continues beyond the displayed region, and red vertical bars indicate contig ends. Each track has its own scale bar (200 kb). In *P.* cf. *toxica*, genes flanking the PKS locus are retained on a single contig, but no PKS gene is present between them. In the azooxanthellate species, the corresponding region is interrupted by contig ends. (**b, c**) Comparison of *P*. sp. sakurajimensis g1143 with the *P. caribaeorum* pseudogene g122a (**b**) and the intact PKS gene g125a (**c**). From top to bottom: domain architecture and exon structure (exons 1–15) of g1143, amino acid identity calculated in 51-aa sliding windows, and the aligned *P. caribaeorum* sequence. Coverage and identity are given relative to g1143 (3,594 aa); note that the y-axis ranges differ between (**b**) and (**c**). In (**b**), red bars mark two premature stop codons in exon 1 of g122a. The nucleotide and amino acid alignments around each stop codon are shown below (TTA [Leu] → TGA at aa 175; CAG [Gln] → TAG at aa 1772), with differences in red. Patched, Patched family domain; KS, ketosynthase; AT, acyltransferase; DH, dehydratase; ER, enoylreductase; KR, ketoreductase; ACP, acyl carrier protein; TE, thioesterase.
